# Cryo-electron microscopy of the f1 filamentous phage reveals a new paradigm in viral infection and assembly

**DOI:** 10.1101/2022.11.03.514279

**Authors:** Rebecca Conners, Rayén Ignacia León-Quezada, Mathew McLaren, Nicholas J Bennett, Bertram Daum, Jasna Rakonjac, Vicki A M Gold

**Author notes:** equal first author contribution (Rebecca Conners, Rayén Ignacia León-Quezada, Mathew McLaren).

## Abstract

Phages are viruses that infect bacteria and dominate every ecosystem on our planet. As well as impacting microbial ecology, physiology and evolution, phages are exploited as tools in molecular biology and biotechnology. This is particularly true for the Ff (f1, fd or M13) phages, which represent a widely distributed group of filamentous viruses. Over nearly five decades, Ff has seen an extraordinary range of applications, including in phage display and nanotechnology. However, the complete structure of the phage capsid and consequently the mechanisms of infection and assembly remain largely mysterious. Using cryo-electron microscopy and a highly efficient system for production of short Ff-derived nanorods, we have determined the first structure of a filamentous virus, including the filament tips. Structure combined with mutagenesis was employed to identify domains of the phage that are important in bacterial attack and for release of new phage progeny. These data allow new models to be proposed for the phage lifecycle and will undoubtedly enable the development of novel biotechnological applications.

## Introduction

Filamentous phages are widely distributed viruses infecting all bacterial genera and some archaea^1^. A number of filamentous phages infecting Gram-negative bacteria have been implicated in virulence, for example horizontal gene transfer of cholera toxin in *Vibrio cholerae*^2^, or biofilm formation in *Pseudomonas aeruginosa*^3^. The phages are approximately one micron long and 6-7 nm wide - visualised as “hair-like” filaments by electron microscopy^4^. A notable feature of the filamentous phage lifecycle is that they replicate and egress without killing their bacterial host. Most extensively studied in this group are the F-specific filamentous phages (Ffs) that include viruses f1, M13 and fd and infect cells by binding to the F-pilus of *E. coli*^5^. The phages are 98.5% identical in their DNA sequence and are used interchangeably. Ffs are also strikingly simple - their single-stranded DNA genome encodes just 11 proteins, of which 5 form the phage capsid. This simplicity, plus their high stability, has facilitated their use in modern biotechnology - for example in phage display, as a vaccine carrier, in tissue engineering, as a high-powered lithium battery, and in phage therapy^6–9^. Despite their extensive downstream applications, the structure of the entire phage capsid assembly remains a mystery^10^.

pVIII is the “major” capsid protein that forms the filamentous body of the phage. The tips of the filament are formed of the “minor” capsid proteins which occur in pairs. pIII and pVI form the “leading” end of the infecting phage, which interacts with the host receptors to initiate entry, whereas pVII and pIX form the “trailing” end^11, 12^ (Fig. 1A).

**Figure 1.**
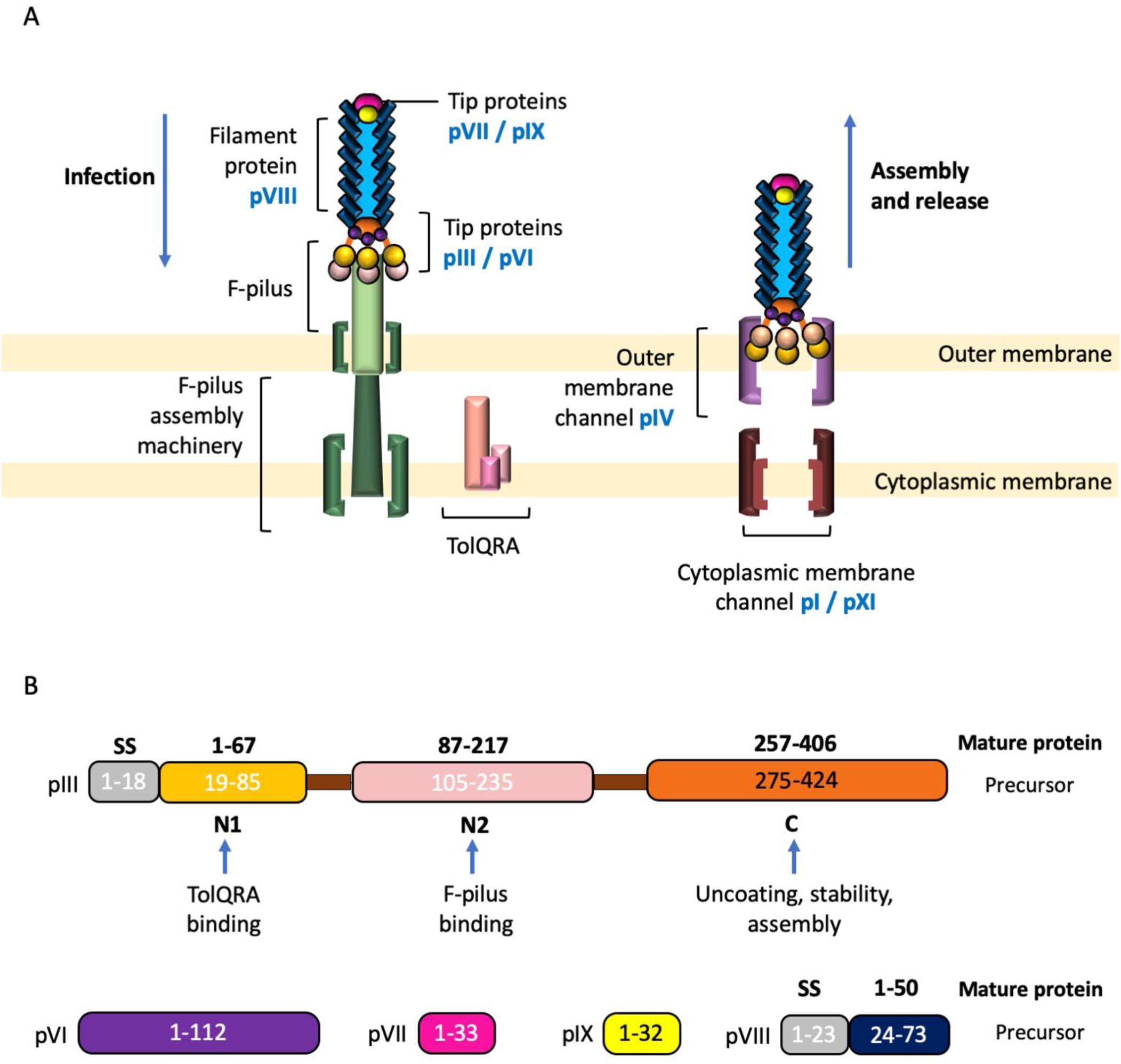
Schematic of phage infection, egress and the proteins that comprise the phage capsid. (A) f1 is comprised of the protein pVIII (blue), which forms the filamentous body of the phage, and the protein pairs pIII (orange/pale pink) and pVI (purple), and pVII (bright pink) and pIX (yellow) that form the tips. For clarity, only three pIII / pVI proteins and one each of pVII / pIX are shown, instead of five of each. pIII binds to the F-pilus (green), which retracts, ultimately allowing phage to reach the TolQRA complex anchored in the cytoplasmic membrane (shades of pink). After DNA injection and genome replication, phage proteins are expressed and assembled. pI / pXI (shades of brown) in the cytoplasmic membrane are dependent on the proton-motive force and ATP hydrolysis by pI. pI / pXI interact with pIV in the outer membrane (mauve), which allows egress across the outer membrane. (B) Schematic to show the f1 capsid proteins and their domains. The numbering for the pIII and pVIII precursor proteins is shown, alongside the numbering for the mature proteins (bold). We have used the numbering for the mature protein throughout the manuscript. SS, signal sequence.

In order for the phage to inject its DNA into the bacterial cytoplasm, it must navigate both the *E. coli* outer and cytoplasmic membranes. Adsorption of the phage to the F-pilus is initiated by the tip protein pIII, which has been divided into 3 domains: N1, N2 and C (Fig. 1B). Specifically, the N2 domain binds to the F-pilus tip^13^ (Fig. 1A), which has two important effects: 1) the F-pilus retracts, bringing the phage to the bacterial surface; and 2) a conformational change occurs in pIII^14, 15^. This frees the N1 domain, enabling it to bind to the periplasmic domain of the secondary receptor TolA, which is part of the TolQRA complex embedded in the cytoplasmic membrane^16^ (Fig. 1A). The mechanism by which phage is able to cross the two membranes is unknown.

The C domain of pIII is involved in virion uncoating and DNA entry into the host cell cytoplasm^17^ (Fig. 1B), where the major capsid protein pVIII is stripped off and integrates into the cytoplasmic membrane^18^. The host transcription and translation machinery replicates the Ff genome, and phage-encoded proteins are synthesised.

Newly synthesised Ff particles exit the cell through a phage-encoded trans-membrane egress machinery, comprised of proteins pI and pXI in the inner membrane, and pIV in the outer membrane (Fig. 1A). We recently determined the structure of pIV by cryo-electron microscopy (cryoEM), revealing how the gates in the channel would need to open to allow phage to egress^19^.

All phage capsid proteins are synthesised initially as integral membrane proteins^20^. The DNA packaging signal is a hairpin, which interacts with the C-terminal residues of the cap proteins pVII and pIX to enable DNA to be incorporated into new virions^21^. The “major” coat protein pVIII is then added, until the entire genome is covered. pIII and pVI are subsequently added as a terminating cap and the phage is released^22, 23^ (Fig. 1A). The phage is now in reverse orientation, with the tip proteins pVII / pIX forming the “leading” end of the egressing filament, and pIII / pVI forming the “trailing” end^11^ (Fig. 1A).

From a structural perspective, the major capsid protein pVIII is the most well-studied to date, with experiments employing fibre-diffraction^24–26^, NMR^27^, and in one case cryoEM^28^. Whilst the latter study provided the first direct view of the pVIII filament, atomic model building was not possible. With improvements in cryoEM methodology^29^, a high-resolution cryoEM structure of a related pVIII protein filament from phage Ike was determined^30^. The four minor cap proteins have not however been amenable to structural characterisation, with the exception of N- terminal fragments of pIII^14, 15, 31^. Therefore, how the capsid proteins interact with one another to form the assembled phage remains elusive.

Currently, cryoEM is the only technique that could be employed to determine a high-resolution structure of an entire filamentous virus. However, determination of protein structures by cryoEM requires averaging thousands of copies of the protein of interest. Due to the length of Ffs (∼1 μm)^4^, the tips are usually present at extremely low abundance in high-magnification images, which makes structural determination challenging. To overcome this, we generated a short version of Ff. This is possible because the length of Ffs can be modulated by changing the size of the packaged DNA^32^. We developed a phage-free plasmid-based system that enables a short version of Ff to be produced based on f1 phage. These are referred to as “nanorods” (Supplementary Fig. 1). This technology has enabled us to determine the first high-resolution structure of a filamentous phage assembly, allowing biochemical data to be reconciled and models proposed for infection, assembly and egress.

## Results

### Generation of the f1-derived nanorods

Nanorods were produced using a two-plasmid system (Supplementary Fig. 1). A nanorod template plasmid was used to generate circular single-stranded DNA 529 nucleotides long, alongside a helper plasmid encoding all f1 phage proteins. On transformation of both plasmids into bacteria, 529-nt circular ssDNA was produced and subsequently packaged into short phage-like nanorod particles by the f1 proteins expressed from the helper plasmid. pVIII contained a Y21M replacement shown previously to result in a more stable conformation of the major coat protein^26^. Nanorods egressed from bacterial cells, and were purified from the medium using CsCl density gradient centrifugation followed by anion exchange chromatography (Supplementary Fig. 2A-C).

### Structure determination of the f1-derived nanorod capsid

Purified nanorods were vitrified by plunge-freezing, and data were collected by cryoEM (Supplementary Table 1). In micrographs, nanorods were observed as structures of ∼80 nm in length (Fig. 2A, Supplementary Fig. 2D), with two distinctive ends: one “round” and one “pointy” (Fig. 2A). Automated particle picking was performed using Warp^33^ for the tips and cryoSPARC^34^ for the filament, and both datasets were subsequently processed using cryoSPARC. After 2D classification of the tips (Fig. 2B-C, Supplementary Fig. 3), particles could clearly be seen to belong to either the round or pointy classes and were sorted accordingly. Additional 2D classification of the filament (Fig. 2D) allowed three different cryoEM maps of the phage to be obtained from a single dataset. All structures had 5-fold symmetry, with the central, filamentous region having additional helical symmetry. The helical parameters were determined using cryoSPARC and were refined to a helical twist of 37.437 ° and a helical rise of 16.599 Å. The final 3D reconstructions, with C5 symmetry applied (and helical symmetry for the central region), produced maps with resolutions of 2.97 Å, 2.58 Å and 2.81 Å for the pointy, filamentous and round structures respectively (Supplementary Fig. 4), into which atomic models were built (Fig. 2E-G, Supplementary movies 1-3).

**Figure 2.**
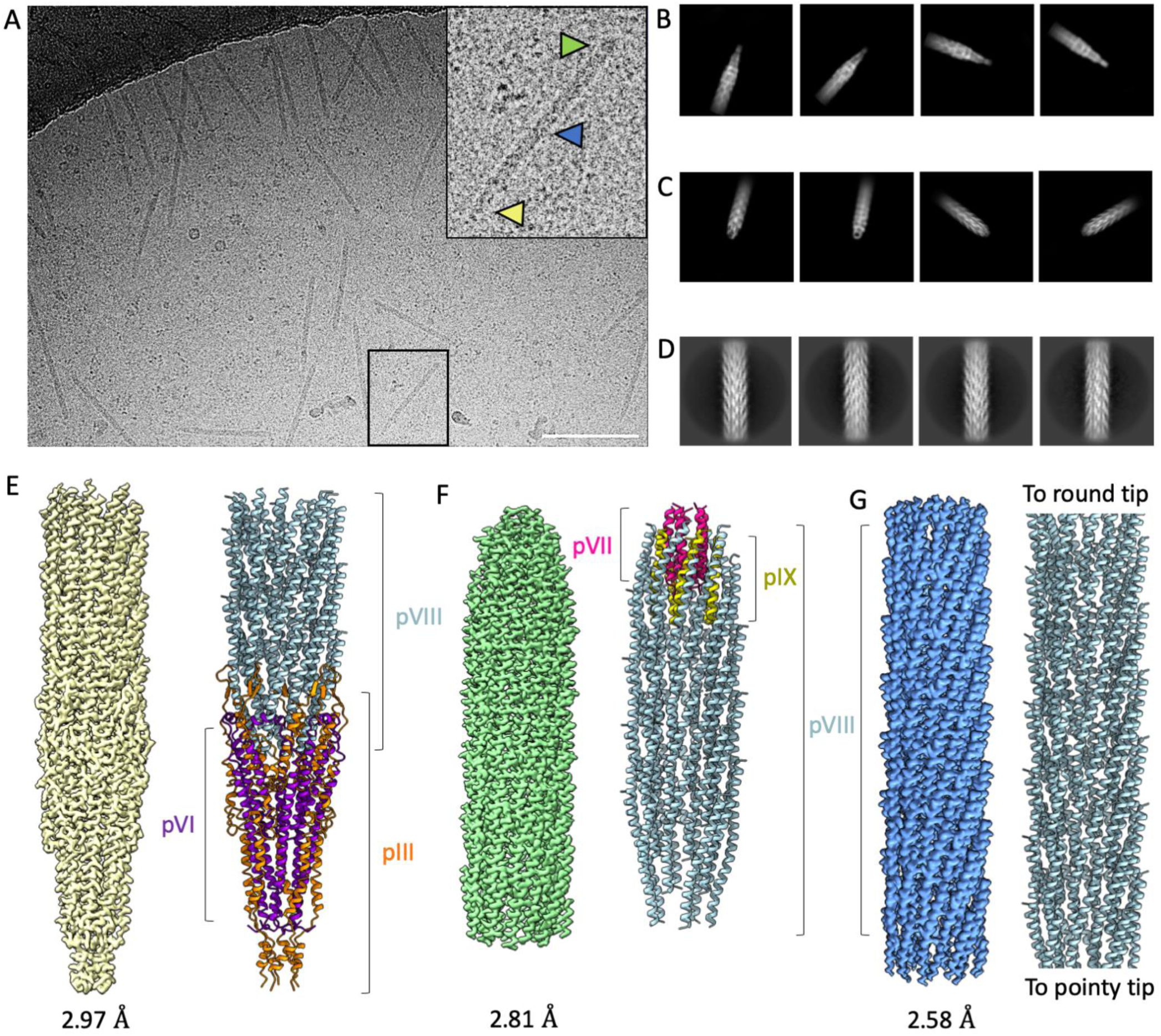
Determination of phage capsid protein structures. (A) Electron micrograph of nanorods showing their pointy (highlighted with yellow arrow) and round (green arrow) tips. The central part of the virion is shown with the blue arrow. Scale bar, 100 nm (B-D) Selected 2D class averages from cryoSPARC, showing (B) four pointy classes, (C) four round classes and (D) four classes of the central, filamentous region. (E) Final 3D reconstruction of pointy tip (yellow) and ribbon diagram with pIII shown in orange, pVI in purple and pVIII in light blue. (F) Final 3D reconstruction of round tip (green) and ribbon diagram with pVII shown in pink, pIX in yellow and pVIII in light blue. (G) Final 3D reconstruction of the central, filamentous region (blue) and ribbon diagram with protein pVIII shown in light blue.

### Structure of the pIII / pVI pointy tip

The pointy tip is comprised of proteins pIII and pVI, present in 5 copies each (Fig. 2E, Fig. 3). pIII is the largest and most structurally complex capsid protein. It is produced with a signal sequence which is cleaved to leave the mature protein of 406 amino acids^35^. Its three domains (N1, N2 and C; of 67, 131 and 150 residues respectively) are separated by two flexible glycine-rich linkers (Fig. 1B). In our structure, the N1 and N2 domains (and the glycine linkers) are not visible, but density was seen for residues 257-404 of the C domain (accounting for the entire C domain minus the final two residues). The pIII C domain is comprised of a number of α-helices of differing lengths (Fig. 3A). Two β-strands form a hairpin loop which extends over the exterior of the main filamentous part of the virion; reminiscent of sepals protecting a flower bud (Fig. 3A box 1). We observed two cysteine residues (354 and 371) in close enough proximity to form an intrachain disulphide bond in the C domain, and these were located at the start and finish of this hairpin loop, pinning it together and further reinforcing the structural motif. pIII is predicted to contain three additional disulphide bonds; two in the N1 domain (between residues 7 and 36, and 46 and 53) and one in the N2 domain (between residues 188 and 201)^31^. The C-terminus of pIII is buried in the centre of the tip, where the five symmetry copies come together to form a distinct stricture in the virion lumen (Fig. 3A box 2, Supplementary Fig. 5A). The lumen measures 8.4 Å across at this point, compared to 20 Å across at its widest point. We observe hydrogen bonds between neighbouring pIII molecules, pinning the tip together (Supplementary Fig. 5B). Two additional inter-chain hydrogen bonds are made between neighbouring pIII subunits; both are situated near rings of methionine residues at the terminus of the pointy tip (Supplementary Fig. 5C).

**Figure 3.**
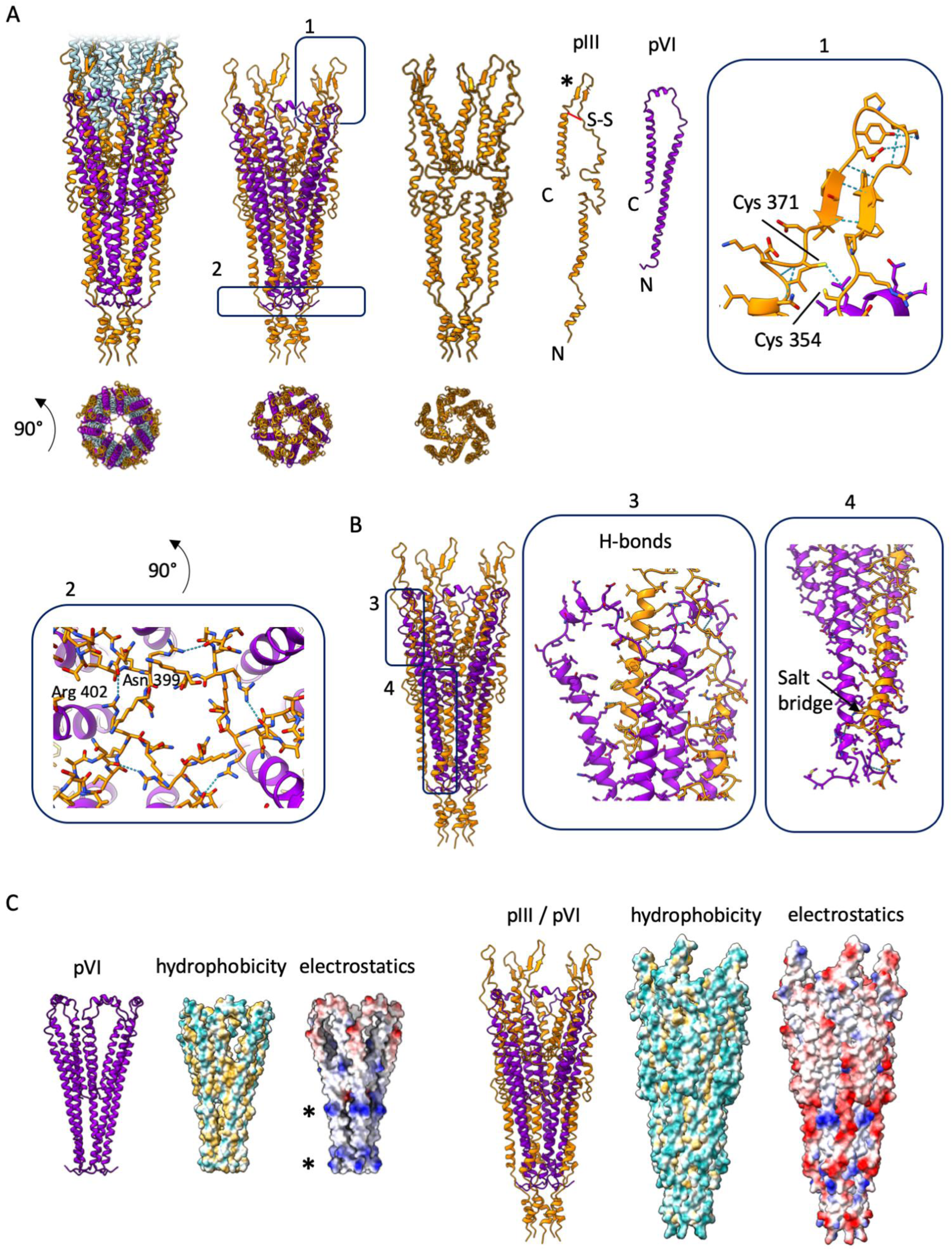
Structure of the pointy tip. (A) Structure of the pointy tip in ribbon representation with pIII shown in orange, pVI in purple and pVIII in light blue. The tip is shown in two views, 90° apart to show a side view and a view looking down the centre of the tip. Proteins pIII and pVI are shown individually as well as in the phage assembled pentameric structure. The disulphide bond in pIII is shown in red and labelled S-S. The * highlights the β-hairpin loop. Parts 1 and 2 are boxed. Part 1 shows hydrogen bonding within the pIII β-hairpin. The cysteine residues are within the correct distance to form a disulphide bond. Part 2 shows a view looking down the phage towards the pointy tip, and a hydrogen bond formed between the sidechain of Arg 402 and the mainchain carbonyl oxygen of Asn 399 from a neighbouring pIII chain. pIII is shown as sticks, pVI as ribbons, hydrogen bonds as light blue dotted lines. (B) Interactions between pIII and pVI. Parts 3 and 4 are boxed, and show the hydrogen bonds (light blue) observed between one pIII molecule and two neighbouring pVI molecules, and the salt bridge (Glu 277 to Arg 12). pIII makes (C) Pointy tip shown with pVI bundle only (left), and pIII / pVI bundle (right) in side view shown as ribbons, and surface views of hydrophobicity and electrostatic charge. The most hydrophilic residues are shown in cyan and the most hydrophobic in mustard yellow; negative residues are shown in red and positive residues in blue. * denotes the rings of positive charge in pVI.

pVI is a 112-residue, mostly hydrophobic protein which lacks a signal sequence (Fig. 1B). Density was observed for the entire pVI protein, which is composed of 3 α-helices arranged in a “U” shape (Fig. 3A). The N-terminus of the protein forms the longest α-helix of 54 residues in length, which turns into a short 10 residue α-helix, and then into the final C-terminal α-helix of 32 residues. The C-terminus of pVI is buried in the centre of the pointy tip. Each pVI chain forms seven hydrogen bonds with neighbouring pVI chains; four with one neighbouring chain and three with the other. These bonds are mostly concentrated in the area at the centre of the virion near the pVI C-termini (Supplementary Fig. 5D).

pIII and pVI can be seen to interact extensively with each other, within a network of closely-packed helices (Fig. 3). Five copies of the protein pair are arranged symmetrically to form the pointy tip with helices from both proteins being intertwined. pIII makes 12 hydrogen bonds and 1 salt bridge with its two neighbouring pVI molecules (Fig. 3B). The outer surface of pVI (which is pIII-facing) is hydrophobic with two distinct rings of positive charge (Fig. 3C, Supplementary Fig. 6, Supplementary Fig. 7). pIII forms a negatively charged scaffold around it, with hydrophobic interactions being shielded at the centre of the assembled phage (Fig. 3C, Supplementary Fig. 6, Supplementary Fig. 7). The pointy tip is overall negatively charged and hydrophilic on the solvent-exposed outside.

At the pointy tip, there were three areas of density that were unaccounted for in the cryoEM map, lying within the central pore of the virion (Supplementary Fig. 8). The first area, at the very tip of the structure, is surrounded by a ring of phenylalanine residues, a lower ring of methionine residues and a third ring of asparagine residues from the pIII protein (Supplementary Fig. 8B). The second, smaller area of density is adjacent to the first, with the asparagine ring defining one end and a second methionine ring the other end (Supplementary Fig. 8B). The third area is located closer to the main body of the filament, where 5 symmetrical curved tubes of density were observed lining a hydrophobic section of the lumen formed by the pVI protein (Supplementary Fig. 8C). These tube-like densities are all consistent with the size and shape of fatty acid molecules, which is supported by the knowledge that lipids are not uncommon in the capsids of many different types of virus^36^. The binding environment provided by the phage corroborates this observation, with hydrophobic residues lining most of the pocket, and hydrophilic residues available to stabilise the acid group of the fatty acid (Supplementary Fig. 8C). Fatty acids were modelled into the map according to their size, suggesting one molecule of octanoic acid (C8) at the tip, followed by one molecule of butanoic acid (C4) and then 5 symmetry-related molecules of dodecanoic acid (C12) in the third site (Supplementary Fig. 8C). Lipids are usually acquired in phages during their assembly, and are thought to aid the infection process by diverse mechanisms^36^.

### Structure of the pVII / pIX round tip

pVII and pIX are small hydrophobic proteins of only 33 and 32 amino acids respectively (Fig. 1B). They both lack a signal sequence, form a single α-helix, and are present in the tip at five copies (Fig. 4A). In comparison to the pointy tip, the rounded one has a much simpler composition. Five copies of pVII form a helical bundle, packed together using mainly hydrophobic interactions (Fig. 4A, Supplementary Fig. 9), as per the pointy tip. Hydrophobic residues line the sides of the helices that pack together, and the interior of the bundle is also predominantly hydrophobic. The terminus of the rounded end of the phage is comprised of the N-terminus of pVII, which has an overall positive charge (Fig. 4A, Supplementary Fig. 10). The helices have typical intrahelical hydrogen bonding, with two interchain hydrogen bonds between neighbouring subunits (Fig. 4B). All proteins are arranged with their C-termini orientated towards the DNA and the main body of the phage. This orientation explains why display of peptide or protein at the round tip of Ff is possible only when linked to the N-termini of pVII and pIX^12^.

**Figure 4.**
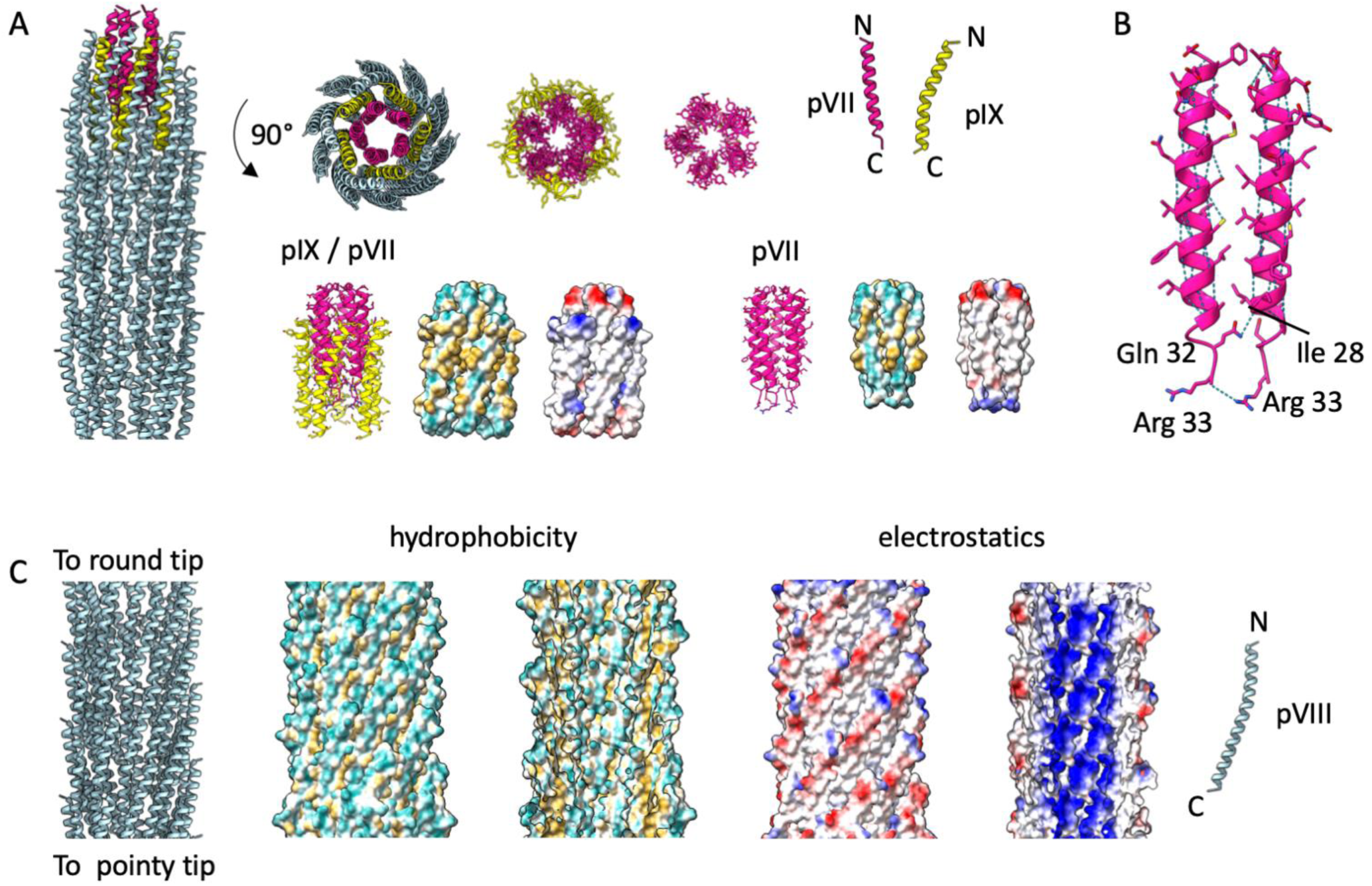
Structure of the round tip and filament. (A) Structure of the round tip in ribbon representation with pVII shown in pink, pIX in yellow and pVIII in light blue. The tip is shown in two views, 90° apart to show a side view and a view looking down the centre of the tip. Proteins pVII and pIX are shown individually, as pentamers, and in the complete structure. Side views are also shown as surfaces coloured by hydrophobicity and electrostatic charge. The most hydrophilic residues are shown in cyan and the most hydrophobic in mustard yellow; negative residues are shown in red and positive residues in blue. (B) Hydrogen bonds in the pVII bundle, shown in ribbon representation. Two interchain hydrogen bonds are formed between neighbouring Gln 32 and Ile 28, and Arg 33 and Arg 33. (C) Structure of the pVIII filament shown as, left to right, ribbon diagram, outer surface coloured by hydrophobicity, inner surface coloured by hydrophobicity, outer surface coloured by electrostatic charge, inner surface coloured by electrostatic charge, and a single protein chain. The most hydrophilic residues are shown in cyan and the most hydrophobic in mustard yellow; negative residues are shown in red and positive residues in blue.

### Structure of the pVIII filament

pVIII is the major capsid protein and forms the main body of the filamentous phage particle. It is produced initially with a signal sequence that is cleaved to leave a single α-helix of 50 residues in length (Fig. 1B), with thousands of these helices packing together in a helical array to coat the length of the viral DNA genome (Fig. 2G). The α-helices are slightly curved; with their C-termini projecting into the phage lumen, resulting in an overall highly positive charge (Fig. 4C, Supplementary Fig. 7D, Supplementary Fig. 10D). The N-terminus of pVIII is splayed outwards towards the exterior of the capsid. In our structure, density was smeared for the four N-terminal residues (1-4), indicating that these are flexible. The proteins are organised with their N-termini pointing towards the round tip, and their C-termini towards the pointy one (Supplementary Fig. 10D). Ff phages belong to the Class I group of filamentous phages (which also include Ike and If1) on the basis of this five-start helical symmetry, as opposed to class II phages such as Pf1 which have a one-start helix^25^. Despite the different symmetries, the overall packing of the pVIII α-helices is the same in both classes.

Overlaying a protomer of our f1-derived nanorod pVIII with a previously determined f1 phage pVIII structure (2C0W; determined by X-ray fibre diffraction^26^) shows that the two are extremely similar (RMSD of 0.976 Å over 39 pruned pairs; RMSD of 1.537 Å across all 46 pairs; Supplementary Fig. 11). However, when comparing the pentamers by aligning at one protomer, small differences can be seen in the placement of the remaining four chains. Ike and the Ff family share 40% sequence identity (Supplementary Fig. 11), with large differences in amino acid properties in some areas. However, overlaying a monomer of pVIII from the Ike filamentous phage (6A7F; determined by cryoEM^30^) shows an even smaller difference between the helices with an RMSD of 0.679 Å (across all 46 pairs). When comparing the pentamers of the f1-derived nanorod pVIII and Ike pVIII, there are again small differences seen in the packing of the individual helices within the pentamer. Interestingly, Ike has a proline at position 30 while all Ff phages have an alanine. This proline mid-way through the helix was proposed to cause the kink observed in the Ike structure^30^, however the kink is still present in the f1-derived nanorod in the absence of the proline.

### Structure of the assembled phage

The Ff phages can be thought of as pentameric building blocks with the first layer closest to the round tip (Fig. 5A). In layer 1, 10 helices of alternating pIX and pVIII line the outside of the pVII bundle (Fig. 5A). Interactions are mainly hydrophobic, with salt bridges between pIX and pVII subunits, and hydrogen bonds between pIX and pVIII subunits, adding further stability to the tip structure (Fig. 5A). Further copies of the main capsid protein pVIII continue to coat the rounded tip in two further layers (layers 2-3), interacting with the pIX helices by mostly hydrophobic interactions.

**Figure 5.**
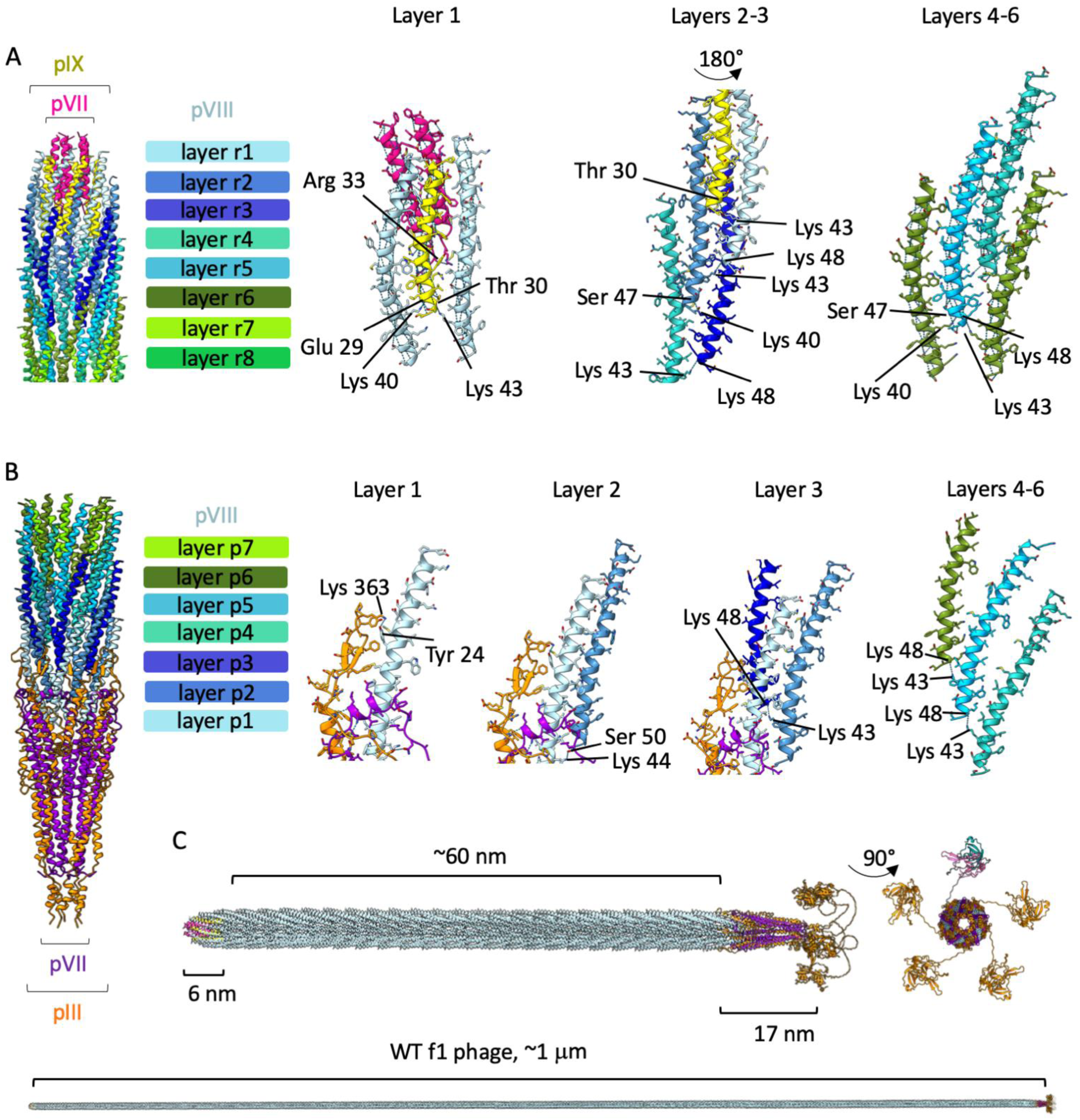
Structure of the assembled phage. Pentameric layers of pVIII are coloured in shades of blue / green according to their position in the filament (e.g. layer r1 in pale blue is the first layer at the round tip, and layer p7 in bright green is the 7^th^ layer at the pointy tip). pVII is shown in pink and pIX in yellow. Hydrogen bonds within the different layers are depicted with blue dashed lines. (A) Left, 10 helices of alternating pIX and pVIII line the outside of the pVII bundle with pentameric layers of pVIII. In layer 1, Interactions are mainly hydrophobic, with salt bridges between pIX and pVII (Glu 29 to Arg 33), and hydrogen bonds between pIX and pVIII (Thr 30 to Lys 43, and Glu 29 to Lys 40). In layers 2-3, pVIII interacts with pIX and neighbouring pVIII molecules via hydrophobic interactions. The pVIII in layer 4 onwards interacts with other pVIII molecules only, again via hydrophobic interactions and with inter-subunit hydrogen bonds between Lys 43 and Lys 48, and Ser 47 and Lys 40 at the C-termini. (B) Left, pentameric layers of pVIII occur at the pointy tip. In layer 1, pVIII interacts with pIII, pVI and neighbouring pVIII molecules, forming hydrogen bonds between Tyr 24 and Lys 363 of pIII, and between Lys 44 and Ser 50 of a neighbouring pVIII from layer 2. In layer 2 pVIII forms hydrogen bonds from Lys 43 to Lys 48 of neighbouring pVIII molecules from layer 3, and in layer 4 onwards, pVIII interacts with other pVIII molecules only, with hydrogen bonds between Lys 43 and Lys 48 from adjacent layers. (C) Composite models of the f1-derived nanorod (top) and WT f1 phage (bottom) were generated by aligning the pointy and round tips with multiple copies of the pVIII filament protein and an Alphafold model of the N1-N2 domains of pIII. The f1-derived nanorod contains ∼230 copies of pVIII, and WT phage ∼2,700 copies. pVII is shown in pink, pIX in yellow, pIII in orange, pVI in purple and pVIII in pale blue. Right, top view of the pointy tip with four of the N1-N2 domains of pIII shown in orange, with one coloured differently to highlight the N1 domain (teal), N2 domain (light pink) and flexible linker regions (grey).

The presence of the major capsid protein pVIII in the round tip means that the few first layers of pVIII will make unique interactions as the round tip section transitions into the filament (Fig. 5A). In layer 1, pVIII interacts with pVII, pIX and neighbouring pVIII molecules. In layers 2 and 3, pVIII interacts with pIX and neighbouring pVIII molecules. The pVIII in layer 4 onwards interacts with other pVIII molecules only. The helical symmetry of the filament is slightly altered as it comes close to the rounded tip, presumably because of the slightly altered binding partners, and the change from helical symmetry of the filament to five-fold symmetry at the very tip. In the filamentous section, pVIII interacts with its neighbours by mainly hydrophobic interactions, with hydrogen bonds observed at the C-termini and mostly involving lysine residues (Fig. 5A).

At the pointy tip, pentameric layers of pVIII interact with the tip proteins pIII and pVI (Fig. 5B). In layer 1, pVIII interacts with pIII, pVI and neighbouring pVIII molecules, with hydrogen bonds linking to layer 2. The hydrogen bonding with pIII occurs in the β-hairpin loop that is stabilised by a disulphide bond (Fig. 3A). In layer 2, pVIII forms additional hydrogen bonds to pVIII molecules from layer 3 (Fig. 5B). In layer 4 onwards, pVIII interacts with other pVIII molecules only, with hydrogen bonds again between adjacent layers.

### Model of the f1 filamentous phage

To visualise the structure of an assembled filamentous phage, we aligned multiple copies of the central, filamentous region with both caps (Fig. 5C). The N1 and N2 domains of pIII, which have been previously observed as small knob-like structures connected to the virion by a “string” of flexible linker^37^, were not visible in our map, and we did not observe any blurred patches of density in our 2D class averages which might have arisen from these domains (Supplementary Fig. 3). Therefore, we used Alphafold^38^ to predict the structure of the N1 and N2 domains, based on the available X-ray structures (1G3P, 2G3P)^14, 31^ (Supplementary Fig. 12A-D). The mostly ß- stranded N1 and N2 domains form a horseshoe arrangement, with the extensive linker regions being unstructured and therefore able to move freely. Although shown symmetrically (Fig. 5C, Supplementary Fig. 13, Supplementary movie 4), these domains could be found in a variety of different orientations surrounding the main body of the pointy tip.

The round tip measures 6 nm from the tip of pVII to the furthest part of pIX, thus comprising only 0.6 % of the total filament (based on a 1 μm total length of WT phage, Fig. 5C). The pointy tip measures 17 nm from the top of the pIII β-hairpin loop to the furthest part of the C1 domain (without N1 and N2), thus comprising 1.6% of the total WT filament (Fig. 5C). The assembled phage displays a clear charge separation. In particular, the N domains and linker regions of pIII are mostly negatively charged, and the phage lumen is overwhelmingly positive due to the C-terminus of pVIII (Supplementary Fig, 10D, Supplementary Fig. 13). Both the phage surface and lumen are lined with hydrophilic residues, with a line of hydrophobic residues mediating interactions between the individual protein subunits (Supplementary Fig. 13).

### Phage DNA

Density was visible for the single-stranded circular DNA genome, however it was not well defined and did not allow a detailed molecular model to be built (Supplementary Fig. 14A, B). At the round end, the C-terminal arginine residues of pVII (Arg 33) and five arginine residues from pIX (Arg 26) form a positively charged ring that butts up to the tip of the DNA (Supplementary Fig. 14C), where the packaging hairpin is expected to be found^21^. The positively charged lumen of the virion is comprised mainly of lysine residues from the C-terminal ends of pVIII molecules (Supplementary Fig. 10D), hence allowing them to interact with the negatively charged DNA molecule. Four lysine residues from each pVIII monomer (Lys 40, 43, 44 and 48) line the DNA cavity (Supplementary Fig. 14D). We fitted a fragment of B-DNA into the density in our map to confirm the approximate dimensions of the nanorod circular ssDNA molecule (Supplementary Fig. 14E).

### The role of pIII in infection

pIII is responsible for receptor binding and subsequent infection of bacteria; our structure allows us to investigate the mechanism. Based on prior functional data, the pIII C domain was divided into C1, C2 and M (transmembrane) sub-domains by mutational analyses^23, 39^. With knowledge of the structure, we now propose that the pIII sub-domain nomenclature follows the position of the α-helices and linkers (Supplementary Fig. 15A).

To examine the roles of specific structural features of the C domain in infection, a series of N1-N2 fusions to truncations of the C domain^40^ were tested for their ability to complement phage containing a complete deletion of gene III (phage f1d3, Supplementary Fig. 15B, C). This analysis showed that phages with short truncations in the C domain (up to 24 residues) were as infective as the wild-type pIII complemented phage (Supplementary Fig. 16, Supplementary Table 2). Mutants with increasingly larger C domain deletions (up to 52 residues) correlated with decreased infectivity by over 30-fold. The mutant with the shortest C-terminal fragment deletion (NdC83; 62 residues) had infectivity equivalent to the negative control phage that contained no pIII (infectivity is six orders of magnitude less than the wild-type pIII-complemented f1d3 phage). We mapped the truncations to the structure of the phage tip, which indicated that the C2 helix (a part of which was previously defined as the Infection Competence Sequence; ICS)^40^ was essential for infection (Supplementary Fig. 16A).

**Figure 6.**
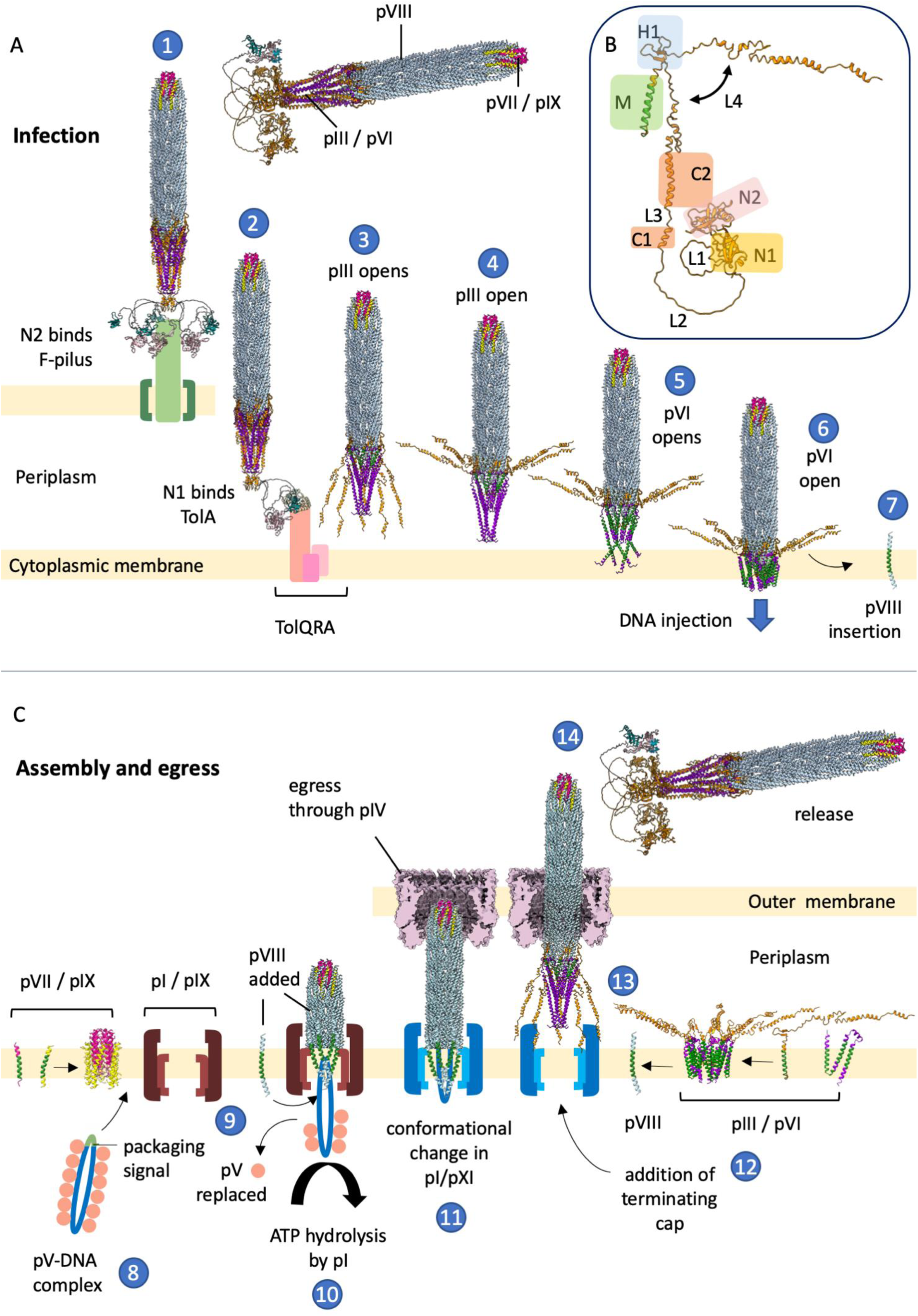
Working model for infection and egress of the Ff family of phages. For clarity, the phage used in the model contains 65 copies of the filament protein pVIII. WT phage contains ∼2,700 copies, so would be ∼42x longer than the phage shown. pVII is shown in pink, pIX in yellow, pIII in orange, pVI in purple and pVIII in pale blue. One of the N1 domains in pIII is highlighted in teal, with its corresponding N2 domain in light pink. (A) Model for infection with (B) domains of pIII highlighted. (1) The N2 domain of pIII binds to the F-pilus and unfolds the N1-N2 hinge to expose the TolA binding site on N1. It is not clear how many N2 domains are needed to bind to the F-pilus for high-efficiency infection. (2) The freed N1 domain of pIII binds to the TolQRA complex. Only one N1-N2 domain is shown for clarity. (3) The C1 and C2 domains of pIII start to move away from the body of the phage at the H1 β-hairpin loop due to flexibility in the L4 linker (B). The N1-N2 domains are not shown for clarity, but could remain bound to TolQRA. (4) Complete opening of the C domain of pIII exposes the hydrophobic pVI pentamer. (5) The exposed hydrophobic regions of pVI insert into the membrane, also allowing insertion of the now exposed pIII transmembrane (M) domain. (6) The pore formed by a pentamer of pIII and a pentamer of pVI allows injection of DNA into the cytoplasm. (7) The first layer of pVIII is exposed on loss of the pointy tip, and proteins move laterally into the bilayer due to opposing charge on the N and C-termini. This reveals the second layer of pVIII, and the process continues. (D) Model for phage assembly and egress with (B) domains of pIII highlighted. (8) ssDNA is packaged by pV, leaving the packaging signal hairpin exposed (green). (9) pV targets the DNA-protein complex to pI / pXI, where pV is replaced by the pVII / pIX round cap. Displacement of pV allows pVIII to diffuse laterally around the DNA. (10) ATP hydrolysis by pI / pXI drives growth of the filament with continuous addition of pVIII. (11) On reaching the terminus of DNA, pI / pXI undergo a conformational change which increases their affinity for the pIII / pVI cap, which form a membrane-bound complex (12). (13) The L4 linker closes around the body of the phage at the H1 β- hairpin loop (B) and the C domain of pIII folds down over the hydrophobic helices of pVI to secede the phage from the cytoplasmic membrane. (14) Phage passes through the pIV secretin complex^19^ in the outer membrane and is released.

Stability of phages can be assessed by investigating their resistance to detergents with different properties. Phages containing wild-type pIII were completely resistant to ionic detergents SDS and Sarkosyl (Supplementary Fig. 17). In contrast, all phages with truncated C domains were completely disassembled in the presence of SDS, and phages with the largest truncations (i.e. that lacked most of C2) were additionally disassembled in the presence of Sarkosyl, a less polar detergent with a larger head group. Interestingly, the predicted octanoic and butanoic acid molecules (Supplementary Fig. 8) would likely be exposed in all of the truncated pIII mutants.

Accessibility of lipids to detergent could explain the change in phage stability; in this case explaining why Sarkosyl (with the larger head group compared to SDS) only affects the stability of the mutants with the largest truncations. This analysis highlights the importance of the C2 domain in infection and stability of phage, and is corroborated by the structure, where the complete C2 helix would shield the hydrophobic pVI core from external solutes (Fig. 3C).

### Membrane integration

All phage capsid proteins integrate into the cytoplasmic membrane upon infection. We therefore investigated their predicted transmembrane regions to understand more about the conformational changes that would need to occur on transitioning from the phage assembled state (Supplementary Fig. 18). Transmembrane sequence analysis with MEMSAT-SVM^41^ predicts that pIII has one transmembrane helix and pVI three (Supplementary Fig. 18A). pVII, pVIII and pIX are all predicted to have one transmembrane helix each (Supplementary Fig. 18A). All capsid proteins are predicted to be orientated with their N-termini in the periplasm and their C-termini in the cytoplasm. Proteins pIII and pVI would need to undergo significant conformational changes to transition from their structure in the phage particle into the membrane-embedded state (Supplementary Fig. 18D). The N-terminal domain of pIII would need to swing out around the β-hairpin loop, and the long ⍺-helix of pVI would need to form two shorter helices. Interestingly, Alphafold predicted that pVI could fold up into a more compact 4-helix form (Supplementary Fig. 18C).

## Discussion

Based on the structure of the f1-derived nanorod, it is possible to reconcile a wealth of phenotypic data and refine current working models for the mechanisms of phage binding, infection, egress and release (Fig. 6).

The pIII tip is responsible for receptor binding on bacteria – first to the tip of the extracellular F-pilus (via the N2 domain)^13^, and second to the TolA protein in the cytoplasmic membrane (via the N1 domain)^42^. The N1 and N2 domains of pIII are bound tightly to each other in a “horseshoe” shape^14, 31^ (Supplementary Fig. 19A). The N2 binding site on the N1 domain overlaps with the TolA binding site^15^ (Supplementary Fig. 19B-C). F-pilus binding must therefore unfold the N1-N2 hinge to expose the TolA binding site on N1^43, 44^ (Supplementary Fig. 19C-D, Supplementary Movie 5). It is not clear how many copies of N2 are required to bind to the F-pilus for high- efficiency infection, but the long glycine-rich linkers between the pIII C-domain and N1-N2 could plausibly allow up to 5 molecules of pIII to clamp around the filament (Supplementary Fig. 19D, Fig. 6A).

Close inspection of the F-pilus machinery reveals an intriguing similarity between the stalk structure (TrwJ), suggested to locate at the tip of the assembled F-pilus^45^, and the pIII C-domain at the tip of f1 (Supplementary Fig. 20A). Both are comprised of pentamers of α-helices, arranged symmetrically, with the same diameter across. When f1 binds to the tip of the F-pilus, these two structures would be directly opposed. Supporting the hypothesis that the two proteins could interact is the observation that the two tips have clear opposing charge, which could allow them to bind with high affinity (Supplementary Fig. 20B).

F-pilus retraction brings phage to the bacterial outer membrane^5^. CryoEM of the F-pilus machinery *in situ*^46^, and of the isolated outer membrane complex^47^, initially suggested that F-pili are assembled directly at the outer membrane. However, a recent high-resolution structure of the purified F-pilus assembly machinery suggests that the dimensions of the trans-envelope chamber are sufficient to accommodate the pilus^45^. f1 is narrower than the F-pilus (Supplementary Fig. 20C); it is therefore possible that phage cross the bacterial outer membrane by passing through the F-pilus machinery directly (Supplementary Fig. 20D). Intriguingly however, phage can infect cells without F-pili in the presence of Ca^2+^ ions, albeit at 6 orders of magnitude reduced efficiency^48^. It therefore cannot be excluded that the role of binding to the F-pilus is to concentrate phage at the outer membrane where crossing of the bilayer occurs by an unknown mechanism. Once the phage tip has crossed the outer membrane, the N1 domain of pIII can locate and bind to TolA, pulling the phage into the periplasm (Fig. 6A).

Phage must next navigate the cytoplasmic membrane. No bacterial proteins have been implicated in the process, thus it is likely also mediated by the “leading" pIII tip. It has been suggested that a conformational change occurring on N1 binding to TolA can initiate a loosening of the interactions between pIII helices in the C-terminal domain of the pointy tip^17^. In support of this, we have demonstrated that the C2 domain of pIII is essential for infection and stability of the phage particle. The L2 linker between the pIII C-terminal domain and the N1-N2 domains (Fig. 6B, Supplementary Fig. 15A) would be sufficiently long (up to 14.8 nm) to allow the C2 domain to reach the TolQRA complex in the cytoplasmic membrane, even if the N1-N2 domains remain bound to their receptors, as has been predicted previously^42^. This idea is in agreement with the fact that the distance between the N1-N2 domains and the C-domain must be maintained for high levels of infectivity^49^. It could therefore be envisaged that the pentameric helices of the pIII C domain (most of C1 and C2) are prised apart by binding TolA in the periplasm, so exposing the hydrophobic pVI pentamer (Supplementary movie 6). The flexible L4 linker between the M and C2 domains of pIII could allow for a twisting out of C1 and C2 about the β-hairpin loop (H1) relative to the transmembrane helix (M domain) (Fig. 6B). Without pIII surrounding pVI and stabilising it, the hydrophobic regions of pVI would become exposed, causing pVI to revert to its 4-helix form and insert into the membrane (Fig. 6A, Supplementary movie 6). The loss of extended pVI would alter the binding partners of the remainder of pIII (up to the β-hairpin structure), now revealing the C-terminal M domain of pIII. This idea is consistent with the fact that pIII has been shown to form a pore in liposomes^50^, and Alphafold predicts the pVI pentamer, and also a pentameric pIII / pVI complex, as channels with sufficient dimension to accommodate DNA (Supplementary Fig. 21A-C). It is interesting to note that the open state tip complex is reminiscent of the family of pentameric ligand gated ion channels^51^. These are also pentamers of transmembrane α-helices, with an extracellular domain containing a distinctive cysteine-loop motif, like pIII. When a ligand binds to the extracellular domain of the ion channels, a conformational change occurs, causing the helices to move apart^51^. It is therefore plausible that a similar mechanism of channel opening allows a pore of pVI and the M domain of pIII to open, allowing injection of DNA into the cytoplasm (Fig. 6A).

The major capsid protein pVIII is inserted into the cytoplasmic membrane on infection. Transmembrane helices have a tendency to orient with positive residues in the cytoplasm, and negative on the opposing side^52^. The two termini of pVIII have clear opposing charges, with the N-terminus being mostly negative, and the C-terminus highly positive (Supplementary Fig. 10D). After opening of the pointy tip, the pVIII charge distribution would thus aid insertion of each helix into the bilayer, via detachment from the main body of the phage and lateral diffusion into the membrane (Fig. 6A). Finally, the terminal pVII and pIX round cap would reach the cytoplasmic membrane, where they would also diffuse laterally into the bilayer due to the presence of a hydrophobic segment each (Fig. 6A).

All of the phage capsid proteins are embedded in the cytoplasmic membrane prior to assembly. It is therefore likely that the conformational changes needed for assembly of proteins into the phage will be similar to the reverse of those occurring during infection. Phage assembly requires energy provided by the pI component of the pI / pXI complex (an ATPase), and a proton motive force^53^. The DNA first is packaged by a phage encoded protein called pV, which wraps around the DNA in a rod of dimers, leaving the packaging signal hairpin exposed^54^ (Fig. 6C).

pV targets the DNA-protein complex to the pI / pXI ring-shaped multimer in the cytoplasmic membrane^55^ where the packaging signal hairpin is thought to act as a signal for replacement of pV by the pVII / pIX round cap (Fig. 6C). The positively charged pocket observed inside the pVII / pIX cap in our structure (Supplementary Fig. 14C) could feasibly mediate the interaction with the DNA hairpin. This is corroborated by the fact that the C-terminal residues of pVII and pIX have been found to interact with phage DNA^21^. Binding could cause the displacement of pV and allow pVIII to diffuse laterally around the DNA at the cytoplasmic membrane, via the positively charged C- terminus interacting with the negatively charged phosphate groups of DNA^56^. ATP hydrolysis by pI / pXI would subsequently drive growth of the filament (Fig. 6C).

Termination occurs by addition of the pIII / pVI pointy cap; both of these two proteins are required for the release of filamentous phage from the infected cells^22^. It has been suggested that pI / pXI undergo a conformational change on reaching the terminus of the DNA, which increases the affinity for the cap proteins^57^ (Fig. 6C). Given that our structure shows that the helices of pIII and pVI are tightly intertwined, it is plausible that pIII and pVI form a membrane-bound complex prior to assembly into the phage particle. This is supported by the findings that pIII protects pVI from proteolytic degradation in *E. coli*^22^. It has also been shown that pIII and pVI each associate with the filament protein pVIII prior to assembly into the phage^20^ (Fig. 6C).

The structures of pIII and pVI support a model in which pIII needs to fold over the hydrophobic C-terminal transmembrane helix of pVI in order to secede from the cytoplasmic membrane. The residues of pIII necessary for co-integration of mutants into the virion together with the wild-type pIII^58^ have been mapped onto our structure and can be seen to lie within the transmembrane helix (M), the β-hairpin loop (H1) and the L4 linker (Supplementary Fig. 22). Mutation of the cysteines in the β-hairpin loop of pIII results in phage that cannot assemble^59^, highlighting the importance of stability in the predicted hinge region (Fig. 6B, Supplementary Fig. 22). The residues essential for assembly therefore all lie within a 93-residue stretch corresponding to the C-terminus of pIII including the L4 linker. The C1 region, L3 and C2 region are not essential for assembly. It is plausible that the H1-L4 region is key for the conformational changes that would need to occur, causing the L4 linker region to swing down to the closed state around the hinge. With respect to termination, a pIII C-domain truncation that is 83 residues long cannot terminate assembly, but a fragment 10 residues longer can^23^. Mapping these 10 residues onto our structure locates the key amino acids required for termination in the small loop where C2 joins the L4 linker (Fig. 6B, Supplementary Fig. 22). For the virion to be released from the cells, the M domain of pIII will need to be extracted from the membrane, together with pVI, for assembly into the phage particle. It is plausible that pIII needs to be of a sufficient length to secede the pVI hydrophobic helices from the membrane, disrupting the hydrophobic interactions with the phospholipid bilayer, pVIII, and/or the pI-pXI transmembrane complex, resulting in release.

Mixing different pIII mutants within a virion has been reported to affect entry and release differently. For example, the complete C domain (lacking N1-N2) was shown to complement the assembly deficiency of the NdC83 internal deletion mutant, but not its lack of infectivity^17^. Furthermore, it was shown that NdC83 has a dominant-negative effect on infectivity of the virion when combined with the wild-type pIII^60^. These two observations differentiate triggers and conformational transitions involved in Ff entry and release, which become examinable now that the structure of the pointy tip has been solved. Practically, combinations described above have been the basis for a major improvement in the power of phage-assisted continuous evolution (PACE)^60^.

In phage display, protein domains or peptides are often fused to the N-termini of either the full-length pIII or to the pIII C terminal domain^61^. However it has been shown that peptides can be linked to the C-terminus of pIII, which we visualise as buried in our structure. Space limitations in the pIII stricture would explain why only an extra 9 residues are tolerated in this position^23^. Longer peptides fused to the pIII C-terminus can only be displayed on the surface of the phage if combined with the wild-type pIII in the same virion, and if a glycine linker is used^62^. In this case the linker must pass between helices in the tip, to allow the peptide to become exposed on the exterior of the phage particle.

### Concluding paragraph

Near-atomic resolution structures of filamentous phage tips in conjunction with structure-function analyses form a new paradigm in virology. We reveal how an intertwined network of α-helices form an extremely stable filament, culminating with a pentameric bundle at the leading tip of the infecting phage. This new knowledge allows the mechanism of cellular-attack to be rationalised, and can now be exploited for mechanistic understanding of the infection and assembly/egress of all filamentous phages. Minor Ff capsid proteins demonstrate great plasticity with respect to incorporation of truncated and mutated subunits into the virions, as well as tolerance to protein fusions of a broad size range^61, 63^. Structural details of the tips will undoubtedly enable improvement of phage display by combining modified virion proteins. In addition, the structures will allow expansion of biotechnological and nanotechnological applications by allowing the precise structure-guided design of novel modification points and 3D display structures.

## Methods

### Generation and purification of the f1-derived nanorods

The nanorods containing a 529-nt circular ssDNA backbone were produced using a two-plasmid system. A helper plasmid (pHP1YM; *IR1-B(C143T*); *gVIII4^am^*, pVIII Y21M) encodes all f1 proteins, and the nanorod template plasmid (pBSFpn529) contains a replication cassette that generates the 529-nt circular ssDNA (details in Supplementary Fig. 1). The nanorods were produced from 1 L of pooled double-transformed host cells. After an overnight incubation of the transformed cell culture, nanorods were purified from the supernatant and concentrated by PEG using standard protocols^64^, with the exception of an increased concentration of PEG due to the short length of the nanorods (15% PEG, 0.5 M NaCl). Furthermore, the nanorod solution after the first PEG precipitation was treated by DNAse and RNAse to remove the cell-derived DNA and RNA. Concentrated nanorods were purified by CsCl density gradient centrifugation using standard Ff (M13, f1 or fd) protocols. Further purification was achieved by anion exchange chromatography using a SepFast™ Super Q column (details in Supplementary Fig. 2). pVIII contains a Y21M replacement shown previously to result in identical conformation of all copies of the major coat protein^26^.

Quantification of nanorods was performed by densitometry of the ssDNA. Briefly, nanorods were disassembled by incubation in ¼ volume of SDS-containing buffer (1% SDS, 1x TAE, 5% glycerol, 0.25 % BPB) at 100°C for 20 min. Disassembled nanorods, along with the quantification standard (serial dilutions of a 529-nt ssDNA of known concentrations purified from the nanorods), were analysed by electrophoresis on 1.2% agarose gels in 1x TAE buffer. All quantification samples were loaded in triplicate. The gel was stained in ethidium bromide, de-stained and imaged using a GelDoc^TM^. Images were analysed using ImageJ^65^. A second-order polynomial function was used to fit the standard curve generated from the ssDNA standards and used to determine the number and concentration of nanorods.

### Cryo-electron microscopy Sample preparation

3 μl of 529-nt nanorods (1.01 x 10^15^ particles/ml) were applied to glow-discharged R1.2/1.3 Cu 300 mesh grids (Quantifoil) and frozen on a Mark IV Vitrobot (Thermo Fisher Scientific) with the following conditions: 4 °C, 100 % relative humidity, wait time 5 sec., drain time 0 sec., blot force 0, blot time 4 sec.

### Imaging

Grids were screened using a 200 kV Talos Arctica microscope (Thermo Fisher Scientific) with a K2 Summit direct electron detector at the GW4 Regional Facility for CryoEM in Bristol, UK. A preliminary dataset was also recorded which was used to determine initial helical parameters. Micrographs used for the final structure were collected on a 300 kV Titan Krios microscope (Thermo Fisher Scientific) with a K3 BioQuantum direct electron detector (Gatan) at the Electron Bio-imaging Centre (eBIC) at Diamond Light Source, UK. Data were collected using EPU software (Thermo Fisher Scientific) with a defocus range from -1.3 to –2.5 μm in 0.3 μm increments. The total dose was 40 electrons/Å^2^ at a magnification of 81 kx, corresponding to a pixel size of 1.10 Å (0.55 Å super-resolution). Further details are shown in Supplementary Table 1.

### Data processing

For the tips, motion correction, CTF estimation and particle picking were performed using Warp^33^ with Box2Net trained using manual picking of ∼1,000 particles, using both round and pointy tips. 594,894 particles were picked and then imported into cryoSPARC 3.2.0^34^. Multiple rounds of 2D classification were used to separate the tips, followed by 3D classification. A total of 86,242 particles of the pointy tip and 255,372 particles of the round tip were selected for final refinement. Iterative 3D refinements and CTF refinements were run for both tips, resulting in resolutions of 2.97 Å and 2.81 Å for the pointy and rounded tips respectively (FSC = 0.143).

To process the central filament, cryoSPARC was used for the motion correction and CTF estimation. The filament tracer tool was used to pick the particles, with an inter-particle spacing of ∼29 Å. A total of 13,430,601 particles were picked, extracted and processed using 2D classification. A total of 1,201,228 particles were selected and used for further refinement. Helical parameters were determined using data from the initial Talos dataset. An initial helical refinement with no symmetry or helical parameters was run and used as a reference for a second refinement with C5 symmetry and no helical parameters. From this map, helical parameters were predicted using the symmetry search tool. The resulting rise and twist values predicted were 16.56 Å and 37.41 degrees respectively. Helical refinements were performed on the Krios dataset using these values as a starting point, with CTF refinement and additional 2D classification performed. The final map achieved a resolution of 2.58 Å using 1,139,813 particles, with helical rise and twist values of 16.599 A and 37.437 degrees respectively, and C5 symmetry applied.

### Modelling

The system of amino acid numbering used throughout is for the mature protein sequences of pIII and pVIII (ie the sequences are numbered from 1 after their signal sequences have been removed). A fibre diffraction structure of fd phage pVIII protein (PDB:2C0W)^26^ was first manually placed in the filamentous cryoEM map using UCSF ChimeraX^66^. Proteins pIII and pVI were easily distinguished from each other on the basis of their length, and were initially built into the pointy density map as poly-Ala chains using Coot^67^ and then their sequences assigned manually. Proteins pVII and pIX were harder to distinguish on the basis of their length, they were also initially built as poly-Ala helices in the round map and their sequences assigned manually. For all three structures, bulky residues, glycine residues and unique sequence patterns were used to guide sequence assignment during model

building. Model building and adjustments were performed using Coot, and the monomeric structures refined using Refmac^68^ from the CCPEM suite^69^. 5-fold and helical symmetry were applied using ChimeraX, and the three complete structures refined again with Refmac. Maps were created without symmetry to check the densities modelled as fatty acids were not symmetry-related artefacts. Models were validated using tools in Coot^67^, Molprobity^70^ and the RCSB validation server^71^. DeepEMhancer^72^ was used for denoising and postprocessing of the maps, and these maps used to aid model building. Model measurements were taken, and figures prepared using ChimeraX^66^. A 12mer of B-DNA was created in Coot, and modelled in the density using Coot.

### Protein structure prediction

Alphafold2^38^ was used for protein structure predictions, with monomers and multimers predicted. MEMSAT- SVM^41^ was used to predict transmembrane regions of the phage proteins.

### Functional analysis of the C domain

Methods for production of C domain mutants of phage, testing infectivity and stability, are described in Supplementary methods.

### Data availability

The 3D cryoEM density maps generated in this study have been deposited in the Electron Microscopy Data Bank (EMDB) under accession codes EMD-15831, EMD-15832 and EMD-15833 for pointy, round and central maps respectively. The atomic coordinates have been deposited in the Protein Data Bank (PDB) under accession numbers 8B3O, 8B3P and 8B3Q. The source image data used in this study have been deposited to the Electron Microscopy Public Image Archive (EMPIAR) under accession number [currently unknown].

## Acknowledgements

This work was funded by a Wellcome Trust Seed Award in Science awarded to VG (210363/Z/18/Z), which along with the University of Exeter, supported RC. MM was supported by a BBSRC responsive mode grant awarded to VG (BB/R008639/1). Work at Massey University and Nanophage Technologies was supported by a Callaghan Innovation Grant (BNANO2101), Palmerston North Medical Foundation, School of Natural Sciences and donations by Anne and Bryce Carmine and an Anonymous Donor. BD received funding from the European Research Council (ERC) under the European Union’s Horizon 2020 research and innovation programme (grant agreement No 803894). We acknowledge Diamond Light Source for access and support of the cryoEM facilities at the UK’s national Electron Bio-imaging Centre (eBIC) at Diamond Light Source [under proposal BI25452], funded by the Wellcome Trust, MRC and BBRSC. We acknowledge access and support of the GW4 Facility for High-Resolution Electron Cryo-Microscopy, funded by the Wellcome Trust (202904/Z/16/Z and 206181/Z/17/Z) and BBSRC (BB/R000484/1). We are grateful to Ufuk Borucu of the GW4 Regional Facility for assistance with screening. We are indebted to George P Smith (University of Missouri) and Marjorie Russel (Rockefeller University) for critical reading and comments on the manuscript.

## Author contributions

RC prepared samples for cryoEM, built the atomic models and analysed the structure. MM collected and processed cryoEM data to determine the structures. RILQ constructed the nanorod production system, generated and purified the f1-derived nanorods. NB conducted functional analysis of the C domain mutants. BD interpreted data and provided resources for electron microscopy. JR and VAMG conceptualised the project, designed the research and obtained the funding. VG wrote the manuscript with RC and JR; all authors commented on the manuscript.

## Materials & Correspondence

Correspondence and material requests should be addressed to JR and VAMG.

## Supplementary figures and tables

**Supplementary Figure 1.**
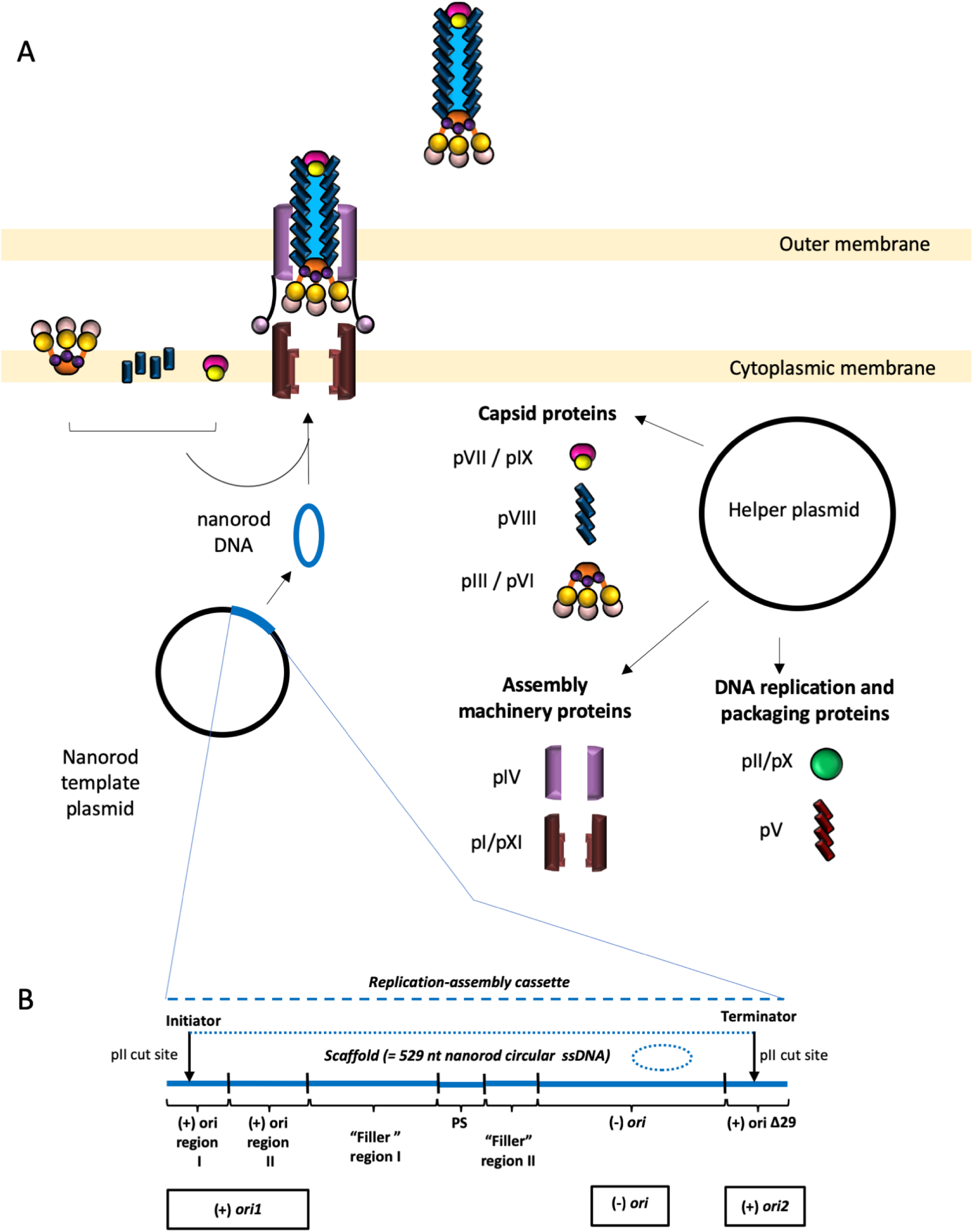
Schematic of the nanorod production system. (A) Schematic drawing of the two-plasmid-based nanorod production system. The system is comprised of a nanorod template plasmid (pBSFpn529) and a helper plasmid (pHP1YM). pBSFpn529 contains a replication- assembly cassette, that in the presence of the helper plasmid (pHP1YM), is replicated to form circular ssDNA of 529 nucleotides, the backbone of the nanorod. The helper plasmid encodes all phage proteins required for replication and assembly of the nanorods. (B) Nanorod replication-assembly cassette within the pBSFpn529 plasmid. “Scaffold” indicates the sequence that is replicated to generate the (+) strand circular ssDNA forming the backbone of the nanorods, between the Ff replicator protein pII cut sites within the replication-assembly cassette. (+) *ori1* corresponds to the complete positive origin of replication, encompassing regions I and II; *(-) ori*, negative strand origin or replication, *(+) ori2*, truncated (+) origin of replication (*Δ29*) encompassing truncated *ori region I* containing deletion of 3’-terminal 29 nt, that serves as the terminator of ssDNA replication. “Filler” indicates DNA segments between the functional sequences. The design of this cassette that includes the *(-) ori* allows replication of both strands once the ssDNA has been excised. This in turn results in an increased nanorod production in comparison to the previously reported nanorod template plasmids containing only the (+) ori that allows only the positive strand replication from the template plasmid^32, 73, 74^.

**Supplementary Figure 2.**
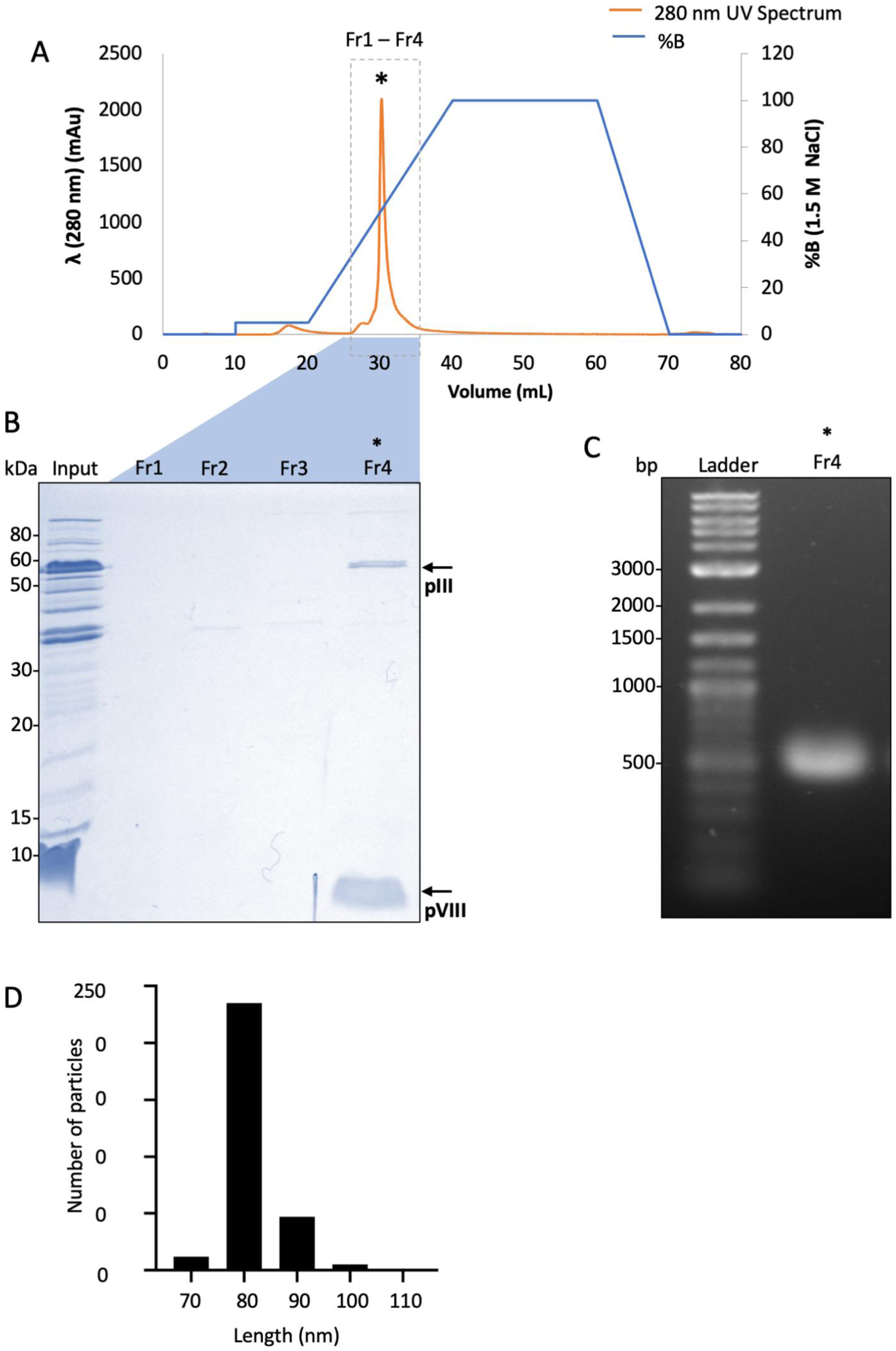
Purification of nanorods by ion-exchange chromatography. Nanorod samples partially purified by CsCl density gradient centrifugation were subjected to chromatography using a SepFast™ Super Q (Strong anion Q, -N+(CH_3_)_3_). (A) Chromatogram depicting UV absorption (mAu; milli Absorption units) at 280 nm (orange line) and %B, corresponding to the salt concentration gradient (blue line, from 0 to 1.5 M NaCl). (B) 16% Tris-glycine SDS-PAGE of the collected fractions. Lanes: kDa, sizes of the molecular weight standard from Novex Sharp pre-stained marker; Input, sample before chromatography (CsCl-purified nanorods); Fr1, Fr2, Fr3 and Fr4, fractions corresponding to the peak indicated in (A). Nanorod proteins pIII and pVIII are indicated by arrows. (C) Fr4 was dialysed and concentrated. To confirm the presence of the nanorods, DNA was disassembled by heating at 100 °C for 10 min, analysed by agarose gel (1.2%) electrophoresis and visualised by staining with ethidium bromide. Lanes: Ladder, 1 Kb plus ladder (a double-stranded liner DNA standard used as a signpost for migration due to the lack of appropriate circular ssDNA standards; numbers indicate sizes of the standard bands in base-pairs, bp); Fr4, the fraction from the ion exchange chromatography (see B). The final nanorod concentration was 2.2 ± 0.2 mg/ml. (D) Histogram of the nanorod length distribution plotted from the length measurements of 300 well-separated particles imaged by negative stain electron microscopy as described in Supplementary methods.

**Supplementary Figure 3.**
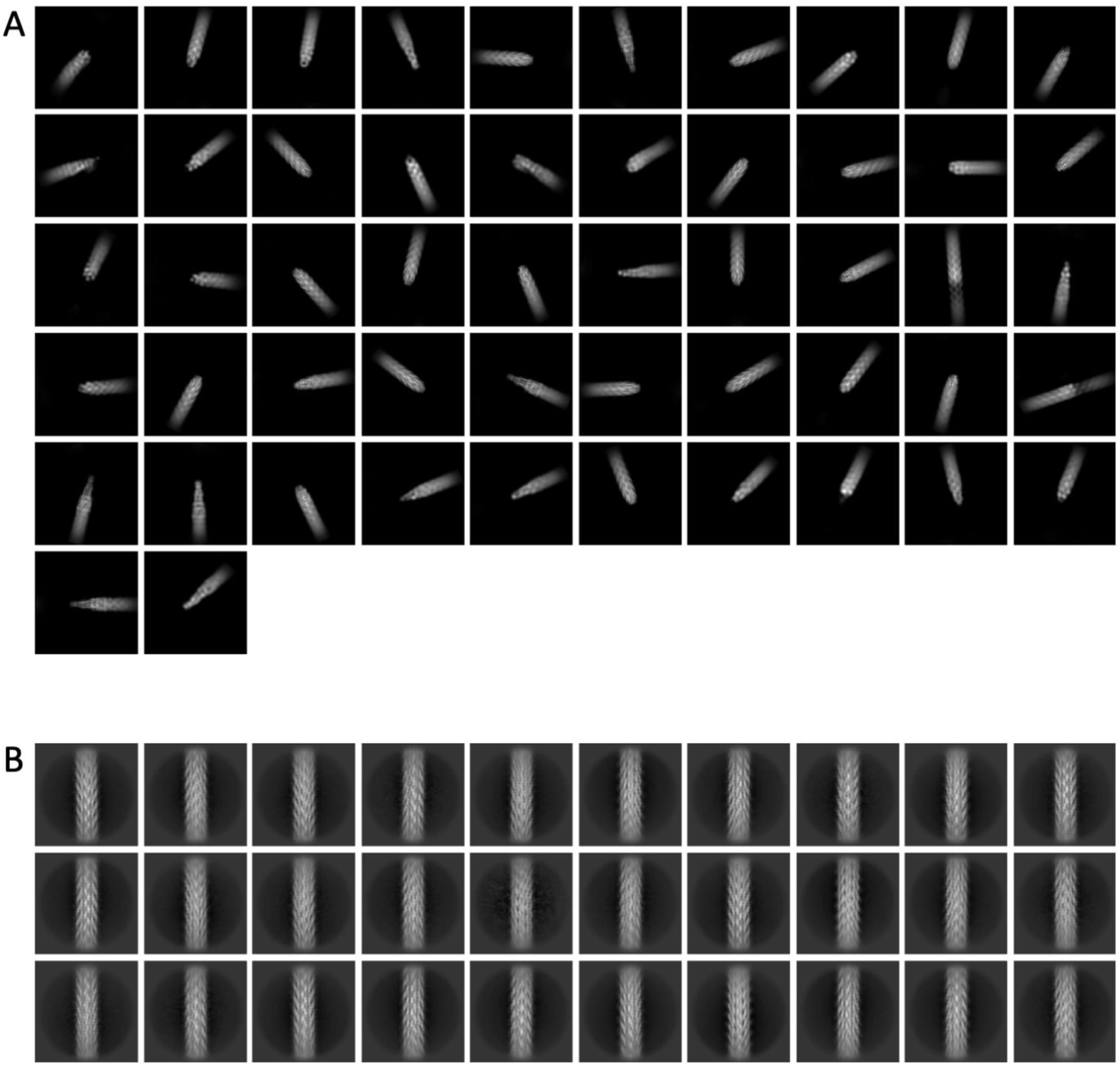
2D class averages. (A) 2D class averages from cryoSPARC for the round and pointy tips. (B) 2D class averages from cryoSPARC for the central, filamentous region.

**Supplementary Figure 4.**
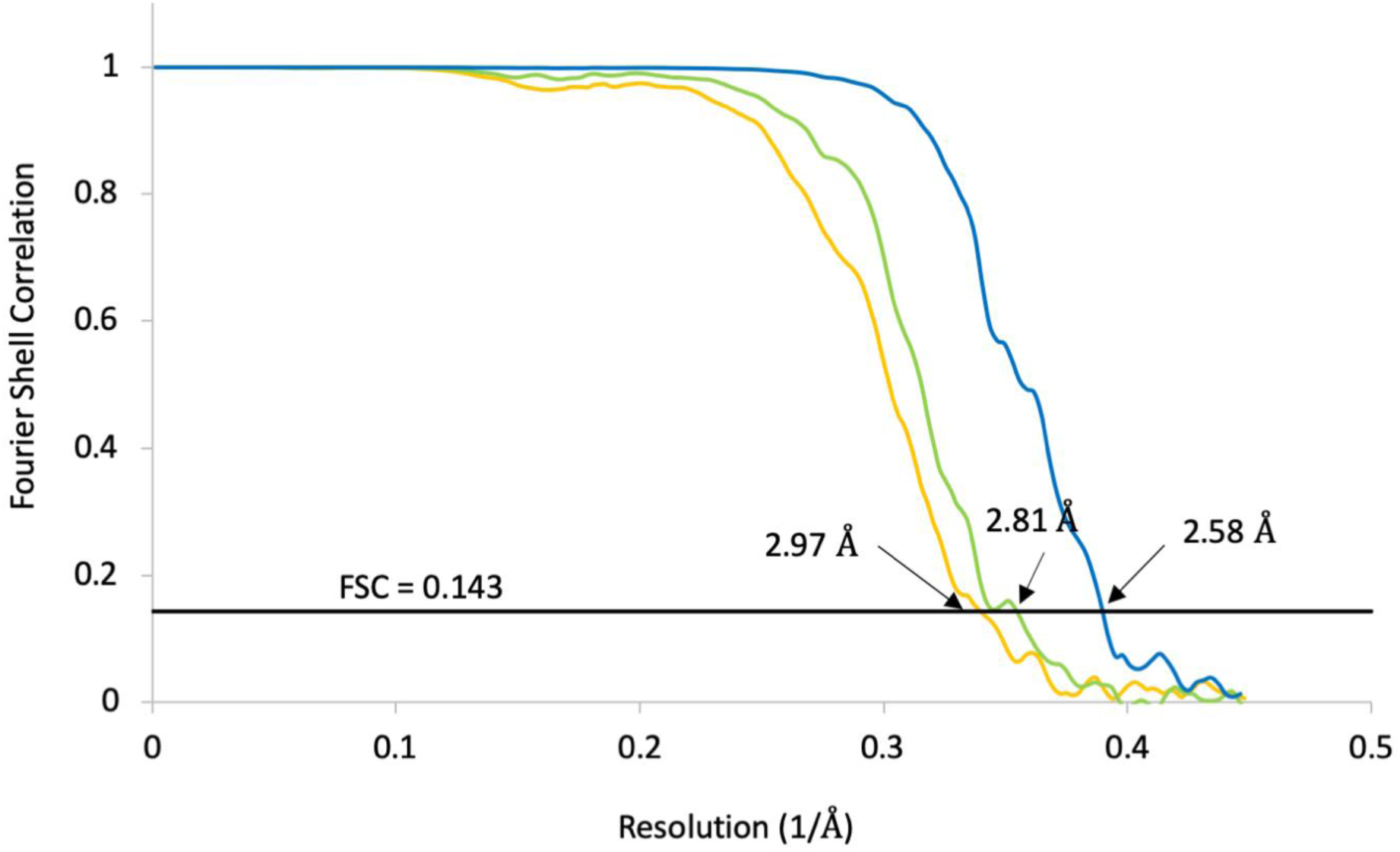
FSC curves. Gold-standard Fourier Shell Correlation (FSC) curves indicate overall resolutions of 2.97 Å, 2.81 Å and 2.58 Å resolution for the pointy tip (yellow), round tip (green) and central part of the nanorod (blue) respectively, at FSC = 0.143.

**Supplementary Figure 5.**
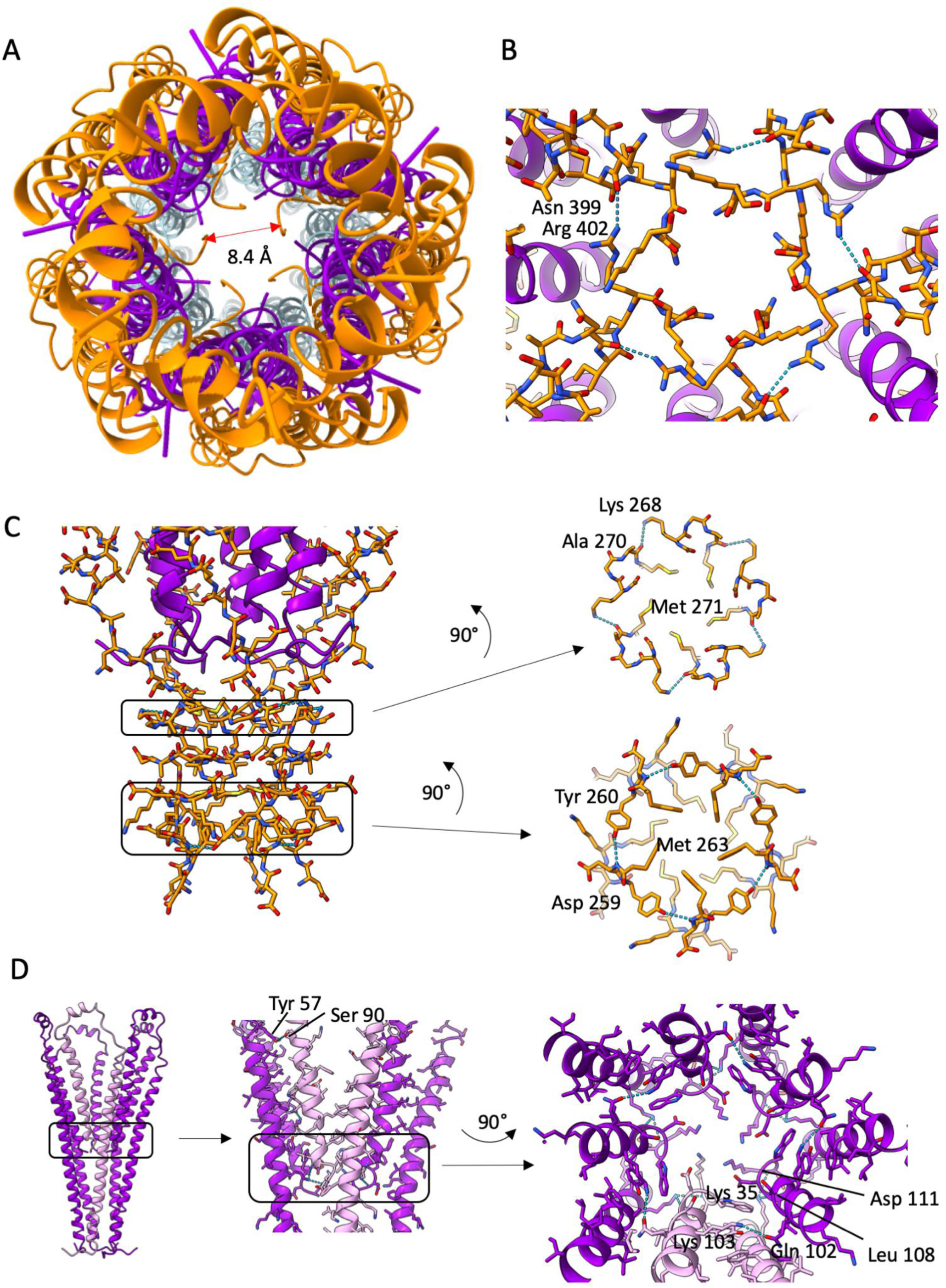
Molecular interactions in pIII and pVI at the pointy tip. (A) The C-terminus of pIII (orange) is buried in the centre of the phage, with the five symmetrical copies forming a stricture of 8.4 Å diameter. pVI is shown in purple. The diameter of the lumen at its widest point in the central region is 25 Å. (B) At the C-terminus of pIII, a hydrogen bond is formed between the sidechain of Arg 402 and the mainchain carbonyl oxygen of Asn 399 from a neighbouring pIII chain. pIII is shown as sticks, pVI as ribbons, hydrogen bonds as blue dotted lines. (C) pIII interchain hydrogen bonds in the pointy tip. Hydrogen bonds are formed between the carbonyl oxygen of Ala 270 and the sidechain of Lys 268 from the neighbouring pIII chain. This hydrogen bonding occurs near a ring of 5 methionine sidechains (Met 271; 1 from each from each pIII in the tip). Moving nearer the tip, hydrogen bonds are formed between the sidechain of Tyr 260 and the backbone amide of Asp 259, again near a motif of a ring of 5 methionine sidechains (Met 263; 1 from each pIII in the tip). (D) pVI pentamer shown in purple, with one chain in lilac. pVI interchain hydrogen bonds are formed between Tyr 57 and Ser 90 of a neighbouring pVI molecule. All remaining interchain pVI hydrogen bonds are found in the region near the C-termini of the protein as indicated in the right panel.

**Supplementary Figure 6.**
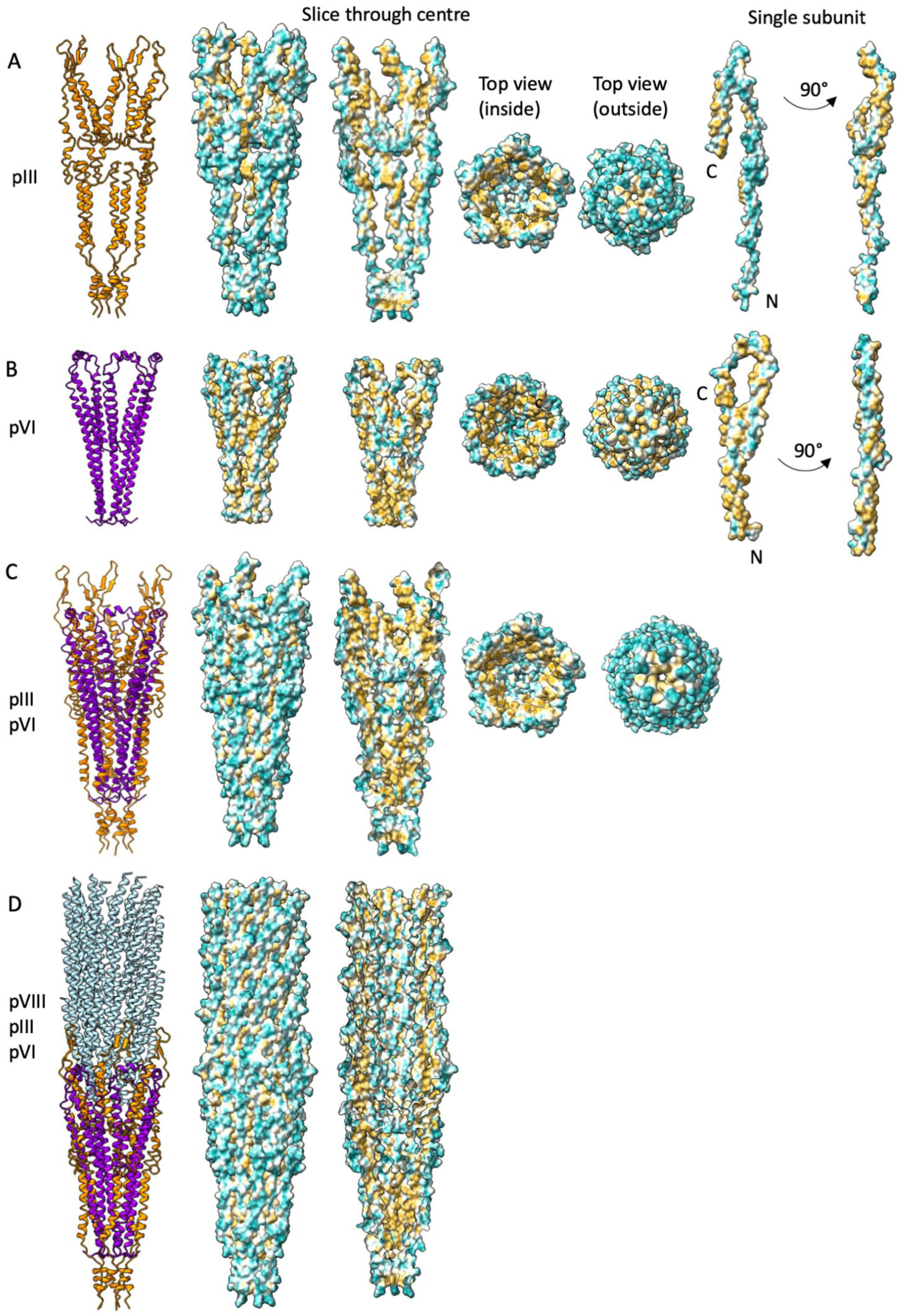
Hydrophobicity of the pointy tip. Surfaces are coloured by hydrophobicity with the most hydrophilic residues shown in cyan and the most hydrophobic in mustard yellow. (A) pIII bundle in side view shown as ribbons, and surfaces in side view, side view sliced through the centre to show the inside of the bundle, top view inside, top view outside. A single chain of pIII is shown in two views 90° apart. (B) pVI bundle in side view shown as ribbons, and surfaces in side view, side view sliced through the centre to show the inside of the bundle, top view inside, top view outside. A single chain of pVI is shown in two views 90° apart. (C) pIII / pVI bundle in side view shown as ribbons, and surfaces in side view, side view sliced through the centre to show the inside of the bundle, top view inside, top view outside. (D) pIII / pVI / pVIII bundle in side view shown as ribbons, and surfaces in side view and side view sliced through the centre to show the inside of the bundle.

**Supplementary Figure 7.**
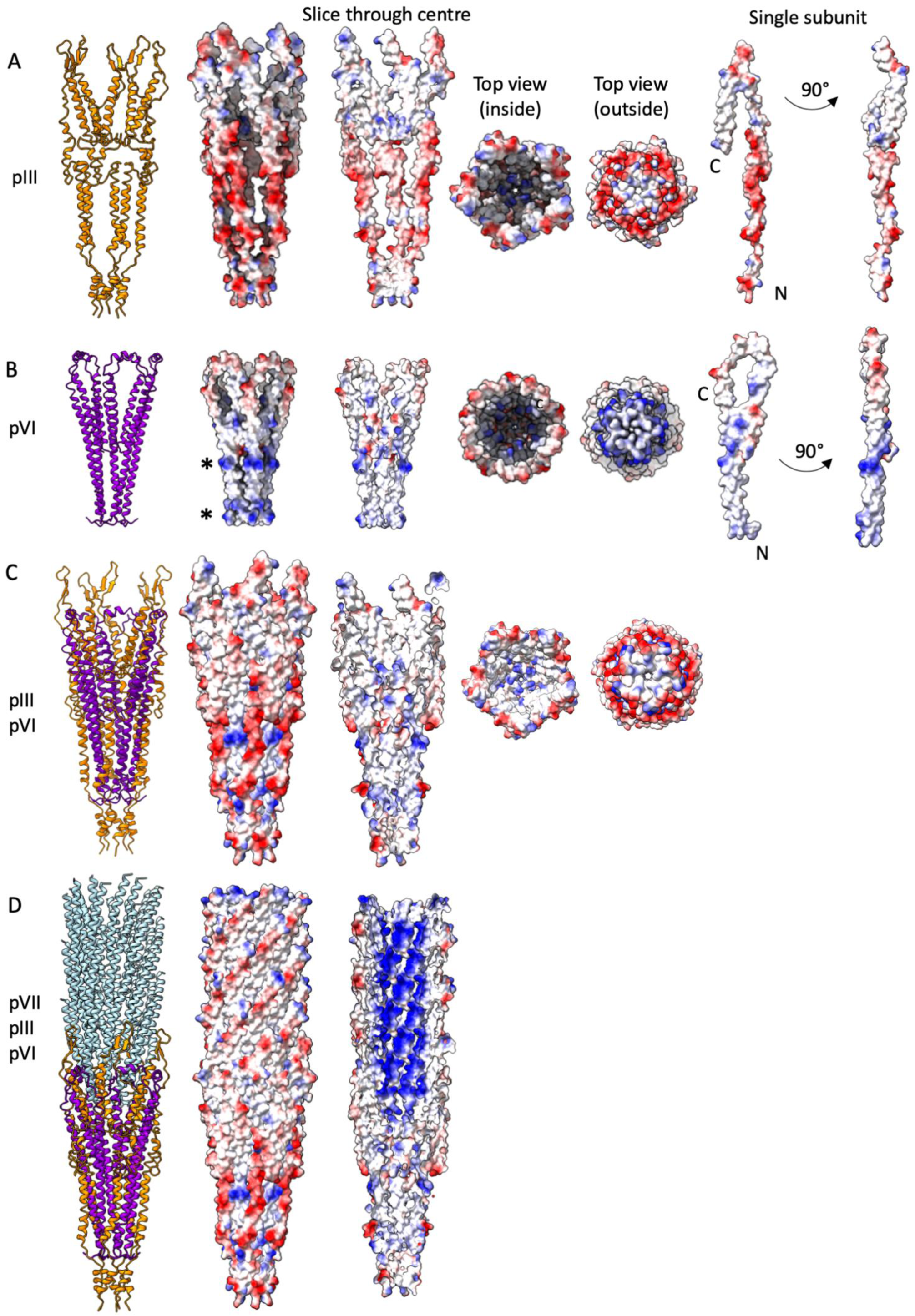
Electrostatics of the pointy tip. Surfaces are coloured by electrostatic surface potential with negative residues shown in red and positive residues in blue. (A) pIII bundle in side view shown as ribbons, and surfaces in side view, side view sliced through the centre to show the inside of the bundle, top view inside, top view outside. A single chain of pIII is shown in two views 90° apart. (B) pVI bundle in side view shown as ribbons, and surfaces in side view, side view sliced through the centre to show the inside of the bundle, top view inside, top view outside. A single chain of pVI is shown in two views 90° apart. * denotes distinct rings of positive charge. (C) pIII / pVI bundle in side view shown as ribbons, and surfaces in side view, side view sliced through the centre to show the inside of the bundle, top view inside, top view outside. (D) pIII / pVI / pVIII bundle in side view shown as ribbons, and surfaces in side view and side view sliced through the centre to show the inside of the bundle.

**Supplementary Figure 8.**
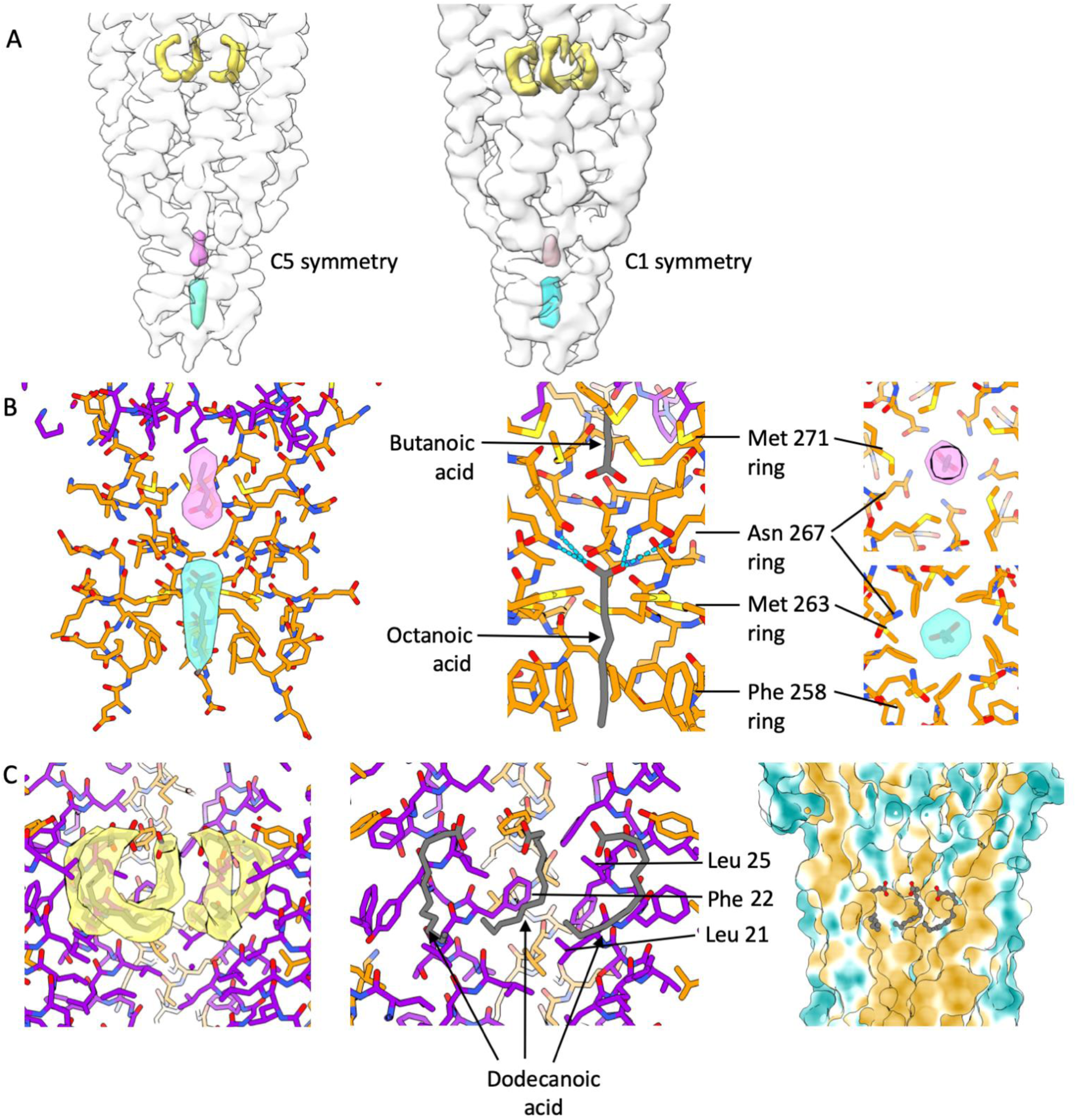
Fatty acid molecules at the pointy tip. (A) Three areas of unaccounted-for density were observed in the central lumen of the pointy tip (coloured cyan, pink and yellow). The three densities were also observed in a map of the pointy tip which was determined with C1 symmetry to rule out symmetry-related artefacts (B) The binding pockets at the tip for the cyan and pink densities are mainly hydrophobic with some hydrogen-bonding possibilities. Therefore we modelled in fatty acids octanoic acid (cyan density) and butanoic acid (pink density). Hydrogen bonds are shown as dotted lines. The binding pocket for the cyan density is composed of rings of phenylalanine, methionine and asparagine residues. The pink density shares the ring of asparagine residues, and has an additional ring of methionines. (C) Five symmetry-related tubes of density observed at the pointy tip had the appearance of fatty acids and were located in a hydrophobic pocket. These have been modelled in as dodecanoic acid (yellow density). The pocket is lined with the hydrophobic residues Leu 21, Phe 22 and Leu 25. Right hand panel shows the virion coloured by hydrophobicity with hydrophobic residues shown in mustard yellow and hydrophilic in cyan.

**Supplementary Figure 9.**
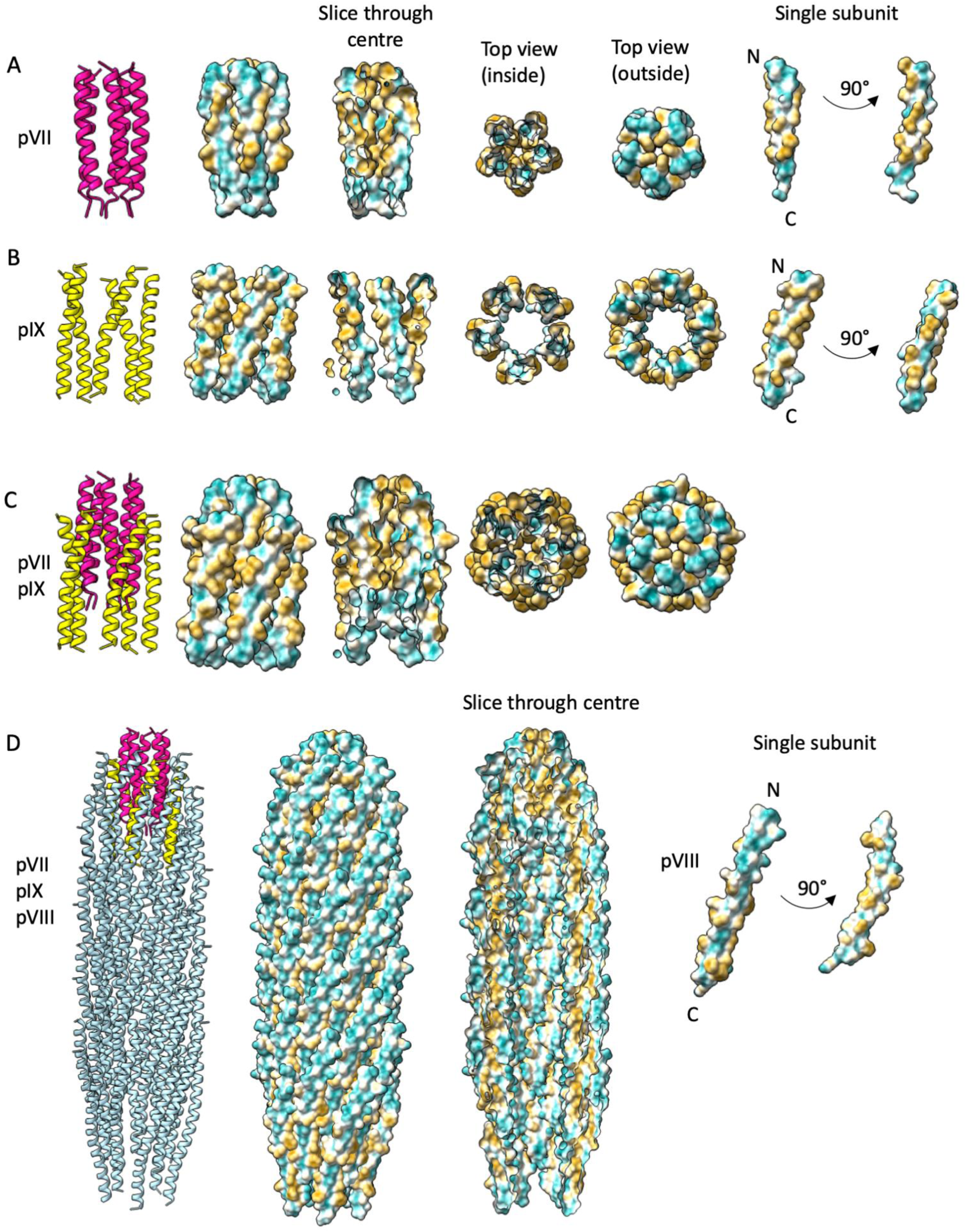
Hydrophobicity of the round tip. Surfaces are coloured by hydrophobicity with the most hydrophilic residues shown in cyan and the most hydrophobic in mustard yellow. (A) pVII bundle in side view shown as ribbons, and surfaces in side view, side view sliced through the centre to show the inside of the bundle, top view inside, top view outside. A single chain of pVII is shown in two views 90° apart. (B) pIX bundle in side view shown as ribbons, and surfaces in side view, side view sliced through the centre to show the inside of the bundle, top view inside, top view outside. A single chain of pIX is shown in two views 90° apart. (C) pVII / pIX bundle in side view shown as ribbons, and surfaces in side view, side view sliced through the centre to show the inside of the bundle, top view inside, top view outside. (D) pVII / pIX / pVIII bundle in side view shown as ribbons, and surfaces in side view and side view sliced through the centre to show the inside of the bundle. A single chain of pVIII is shown in two views 90° apart.

**Supplementary Figure 10.**
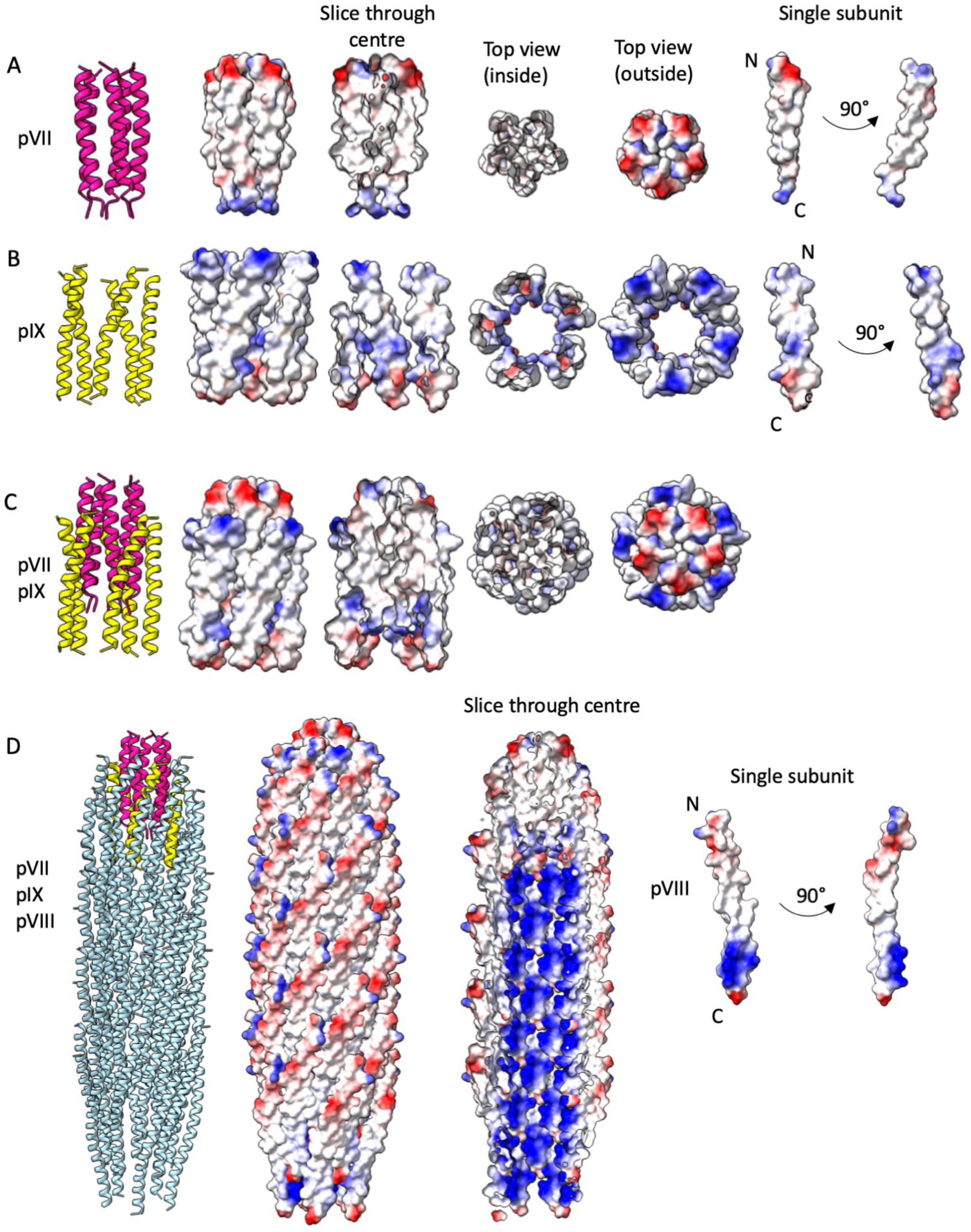
Electrostatics of the round tip. Surfaces are coloured by electrostatic surface potential with negative residues shown in red and positive residues in blue. (A) pVII bundle in side view shown as ribbons, and surfaces in side view, side view sliced through the centre to show the inside of the bundle, top view inside, top view outside. A single chain of pVII is shown in two views 90° apart. (B) pIX bundle in side view shown as ribbons, and surfaces in side view, side view sliced through the centre to show the inside of the bundle, top view inside, top view outside. A single chain of pIX is shown in two views 90° apart. (C) pVII / pIX bundle in side view shown as ribbons, and surfaces in side view, side view sliced through the centre to show the inside of the bundle, top view inside, top view outside. (D) pVII / pIX / pVIII bundle in side view shown as ribbons, and surfaces in side view and side view sliced through the centre to show the inside of the bundle. A single chain of pVIII is shown in two views 90° apart.

**Supplementary Figure 11.**
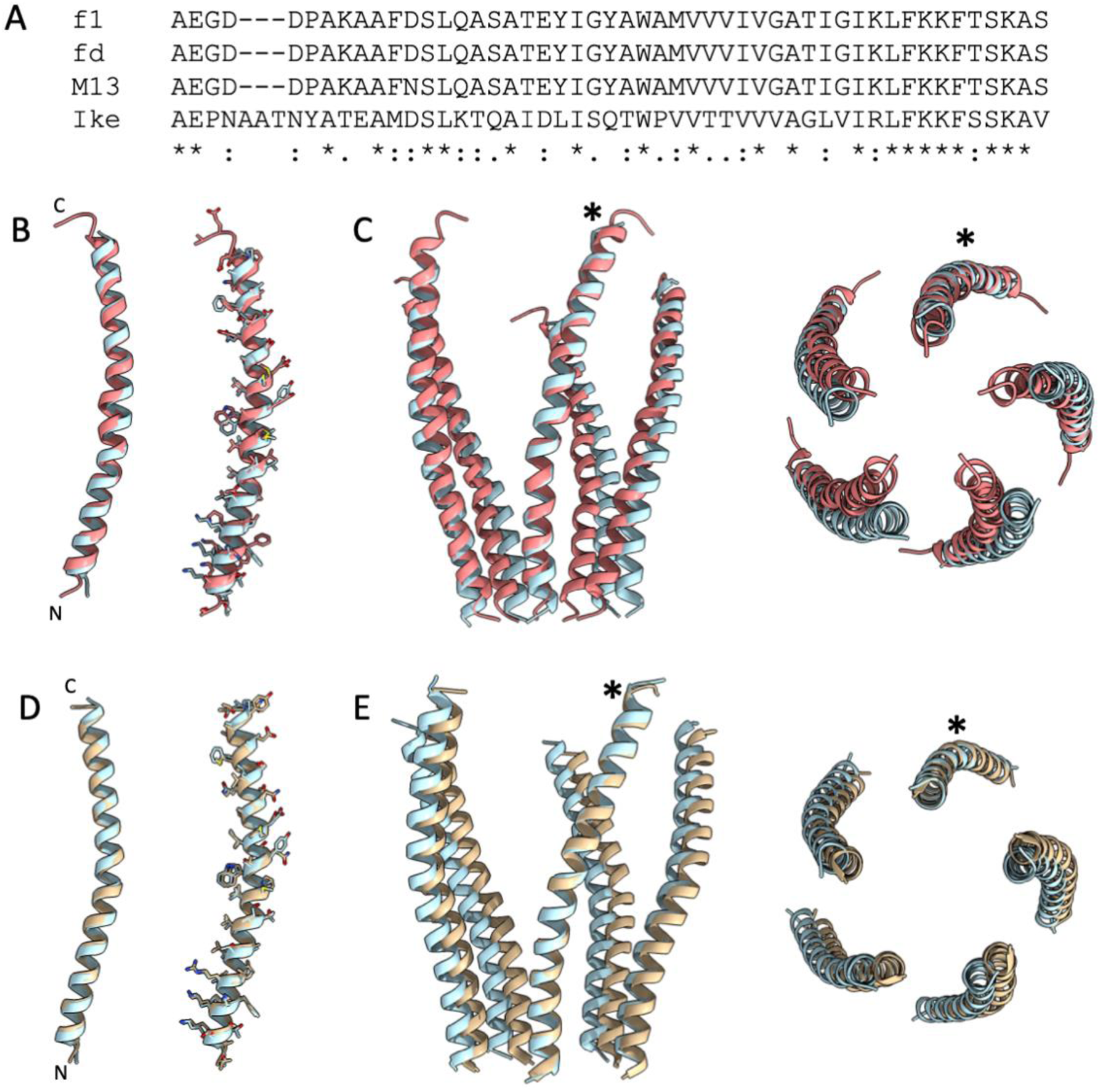
Comparison of the pVIII structure to other phage filaments. (A) Sequence alignment of pVIII from f1, fd, M13 and Ike calculated with T-Coffee^75^. The signal sequences were removed for the alignment. The mature form of pVIII from f1 is 100 % identical to that from fd, 98 % identical to that from M13 (1 amino acid different) and 40 % identical to Ike. Conserved residues are labelled with *, strongly similar with : and weakly similar with . (B) Overlay of the f1-derived nanorod pVIII monomer (light blue) with a monomer of pVIII from fd (2C0W^26^, structure determined by X-ray fibre diffraction, light coral) in cartoon and showing sidechains. (C) Overlay of the nanorod pentamer on the fd (2C0W) pentamer (by aligning one monomer from the nanorod with one monomer from 2C0W) in side and top views. The helices which were aligned are marked with *. (D) Overlay of the nanorod pVIII monomer (light blue) with the Ike monomer (6A7F^30^, structure determined by cryoEM, tan) in cartoon and showing sidechains. (E) Overlay of the nanorod pentamer on the Ike pentamer (by aligning one monomer from the nanorod with one monomer from 6A7F) in side and top views. The helices which were aligned are marked with *.

**Supplementary Figure 12.**
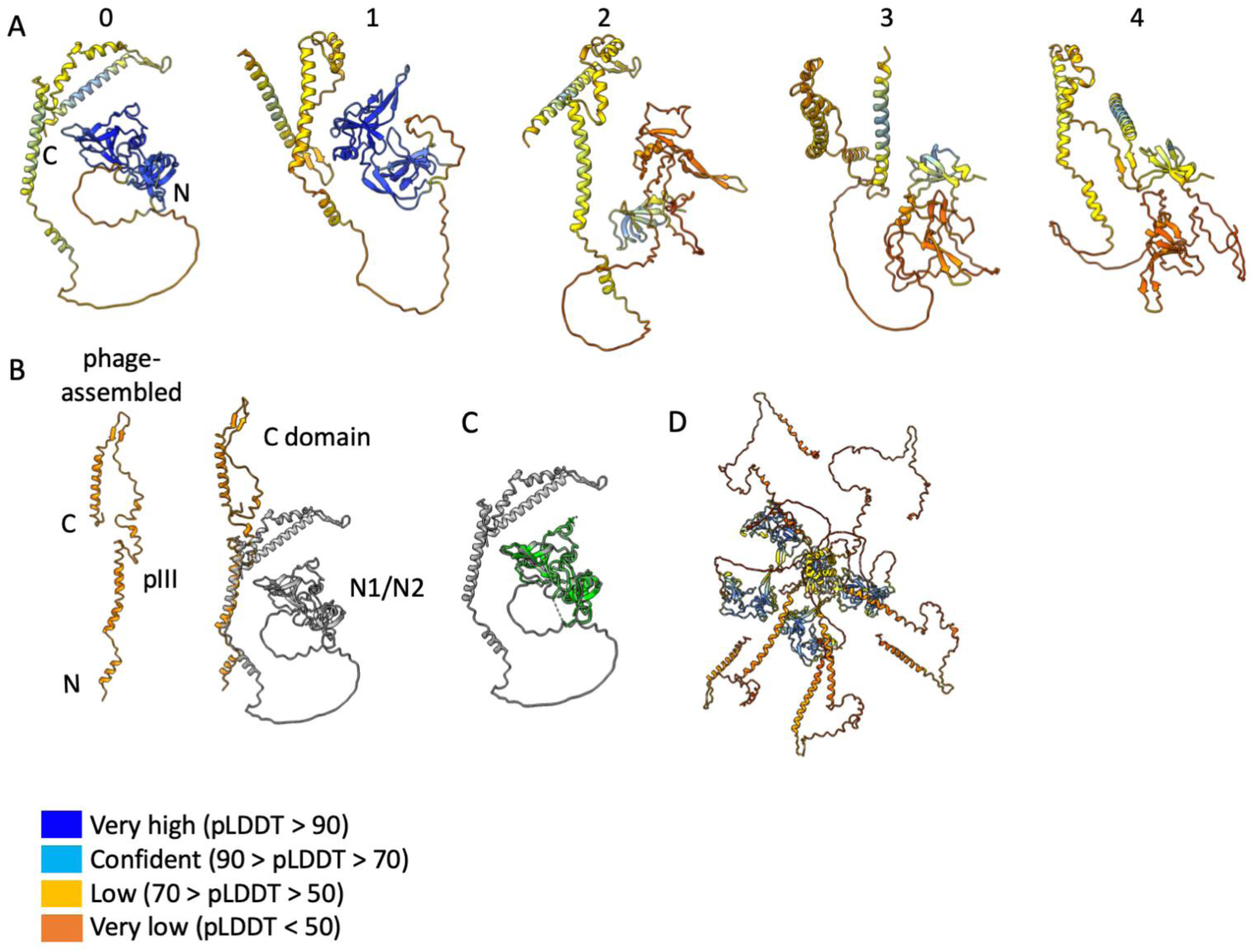
Alphafold predictions of the N1 and N2 domains with linker regions. (A) Alphafold predictions of the full-length pIII monomer coloured by confidence levels with highest confidence in blue, through cornflower blue, then yellow to the lowest confidence in orange. Numbers relate to the confidence ranking from Alphafold (0 is best, 4 is worst). The N and C-termini are labelled in model 0. (B) The top prediction of pIII from Alphafold (model 0 in (A)) shown in grey was overlaid with our structure of the C-terminal domain of pIII shown in dark orange to obtain a model for the relative positions of the N1 and N2 domains. The structure of the pIII C-domain in the phage-assembled structure is shown for comparison. (C) Top prediction of pIII from Alphafold (model 0 in (A)) shown in grey overlaid with the crystal structure of the N1 and N2 domains of pIII (1G3P)^31^ shown in green to demonstrate a good fit (RMSD 0.762 Å between 185 pruned atom pairs). The N1-N2 Alphafold prediction was subsequently modelled onto the tip of the C-domain structure to generate the model of pIII in the assembled phage. (D) Alphafold was not able to predict a reliable model of the full-length pIII pentamer. The top Alphafold prediction of full-length pIII pentamer is shown and coloured by confidence. The N1 and N2 domains are predicted at high confidence, whereas the C domain of pIII is predicted with low confidence, as seen in (A).

**Supplementary Figure 13.**
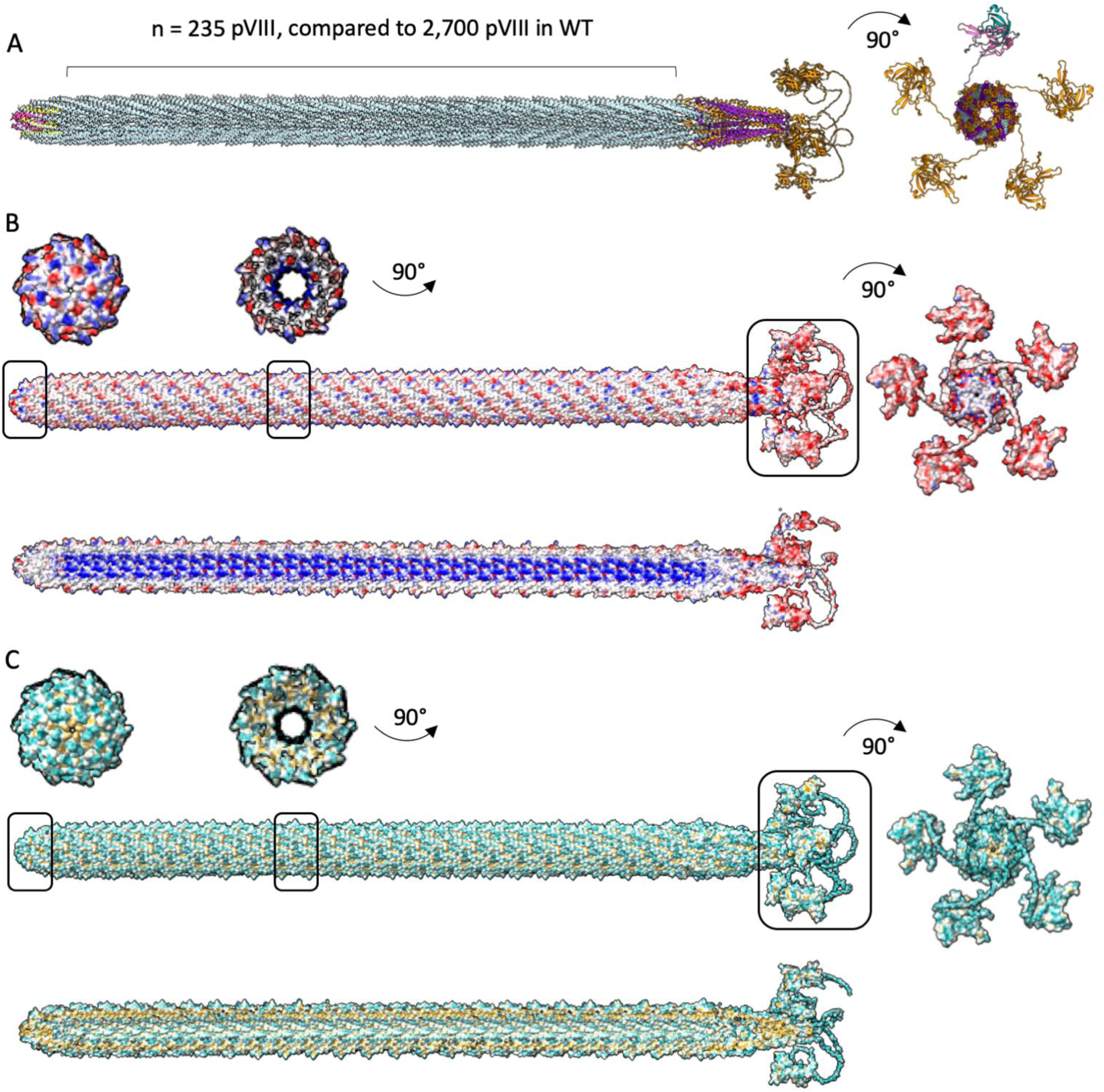
Composite model of the f1-derived nanorod. (A) A composite model was generated by aligning the pointy and round tips with 235 copies of the pVIII filament protein and an Alphafold model of the N1-N2 domains of pIII. WT phage contains ∼2,700 copies of pVIII, so would be 11.5x longer than the structure shown. pVII is shown in pink, pIX in yellow, pIII in orange, pVI in purple and pVIII in pale blue. Right, view of the pointy tip with four of the N-terminal regions of pIII shown in orange, with one coloured differently to highlight the N1 domain (teal), N2 domain (light pink) and flexible linker regions (grey). (B) Phage coloured by electrostatic surface potential. The outer surface and a view sliced through the centre are shown, with negative residues coloured in red and positive residues in blue. Boxed areas show views rotated by 90°. (C) Phage coloured by hydrophobicity. The outer surface and a view sliced through the centre are shown, with the most hydrophilic residues coloured in cyan and the most hydrophobic in mustard yellow. Boxed areas show views rotated by 90°.

**Supplementary Figure 14.**
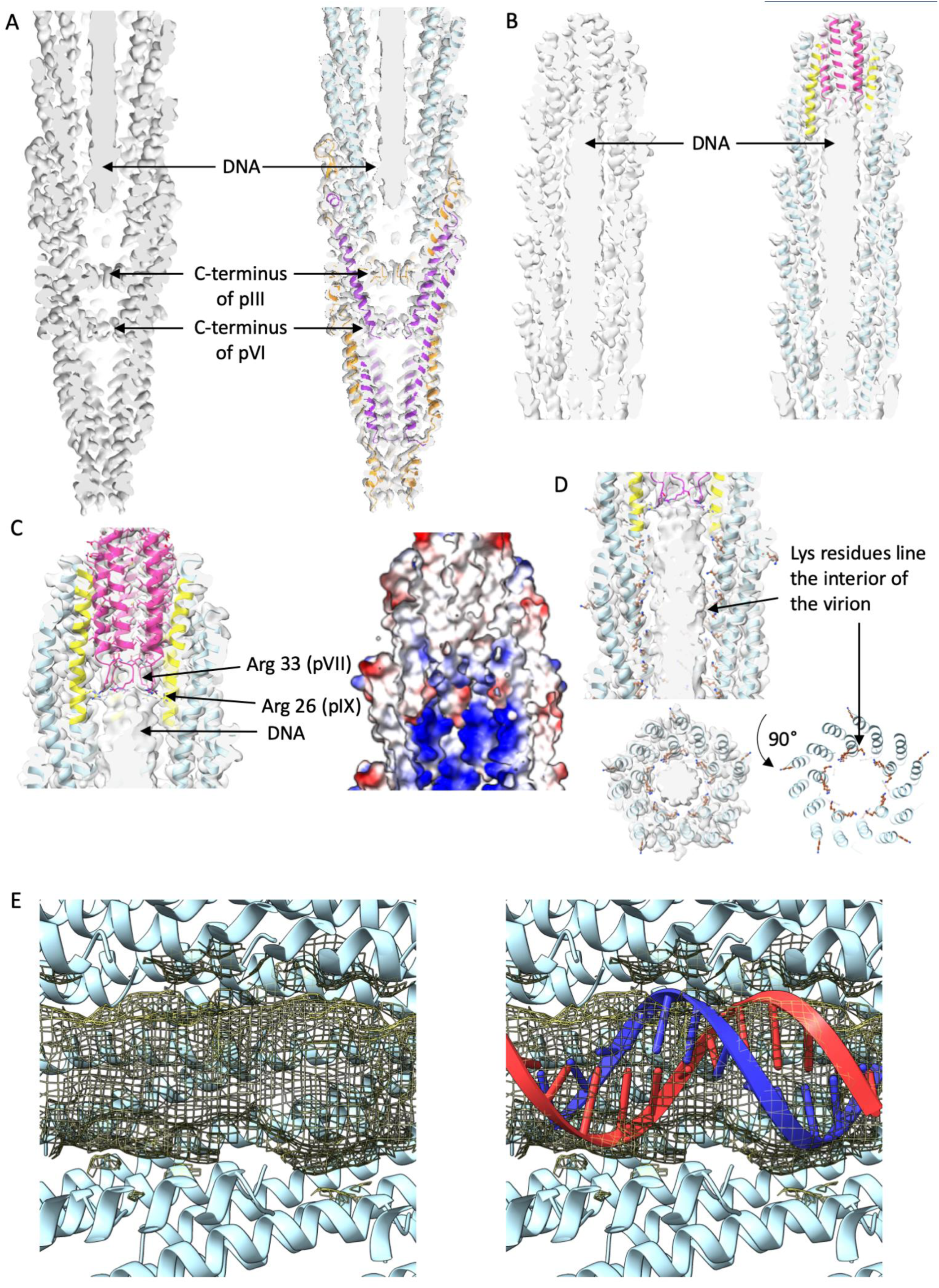
The single stranded DNA genome. (A) A slice through the centre of the pointy tip with density map shown in light grey and protein chains as cartoon. pIII is shown in orange, pVI in purple and pVIII in pale blue. The density for the DNA is clearly visible. (B) A slice through the centre of the round tip with density map shown in light grey and protein chains as cartoon. pVII is shown in pink, pIX in yellow and pVIII in pale blue. The density for the DNA is clearly visible and butts up to the round protein tip. (C) Sidechains Arg 33 of pVII and Arg 26 of pIX are shown as sticks, forming a highly positively charged pocket for the DNA packaging signal hairpin to bind. (D) Lysine residues of protein pVIII are shown as brown sticks. Four lysine residues at the C-terminal end of pVIII line the virion cavity, causing it to be positively charged. Shown in side and top views rotated by 90°. (E) Left; density at the centre of the phage (maps produced with no symmetry applied and displayed around the central axis) and right; with a 12mer of B-DNA modelled (individual strands coloured red and blue).

**Supplementary Figure 15.**
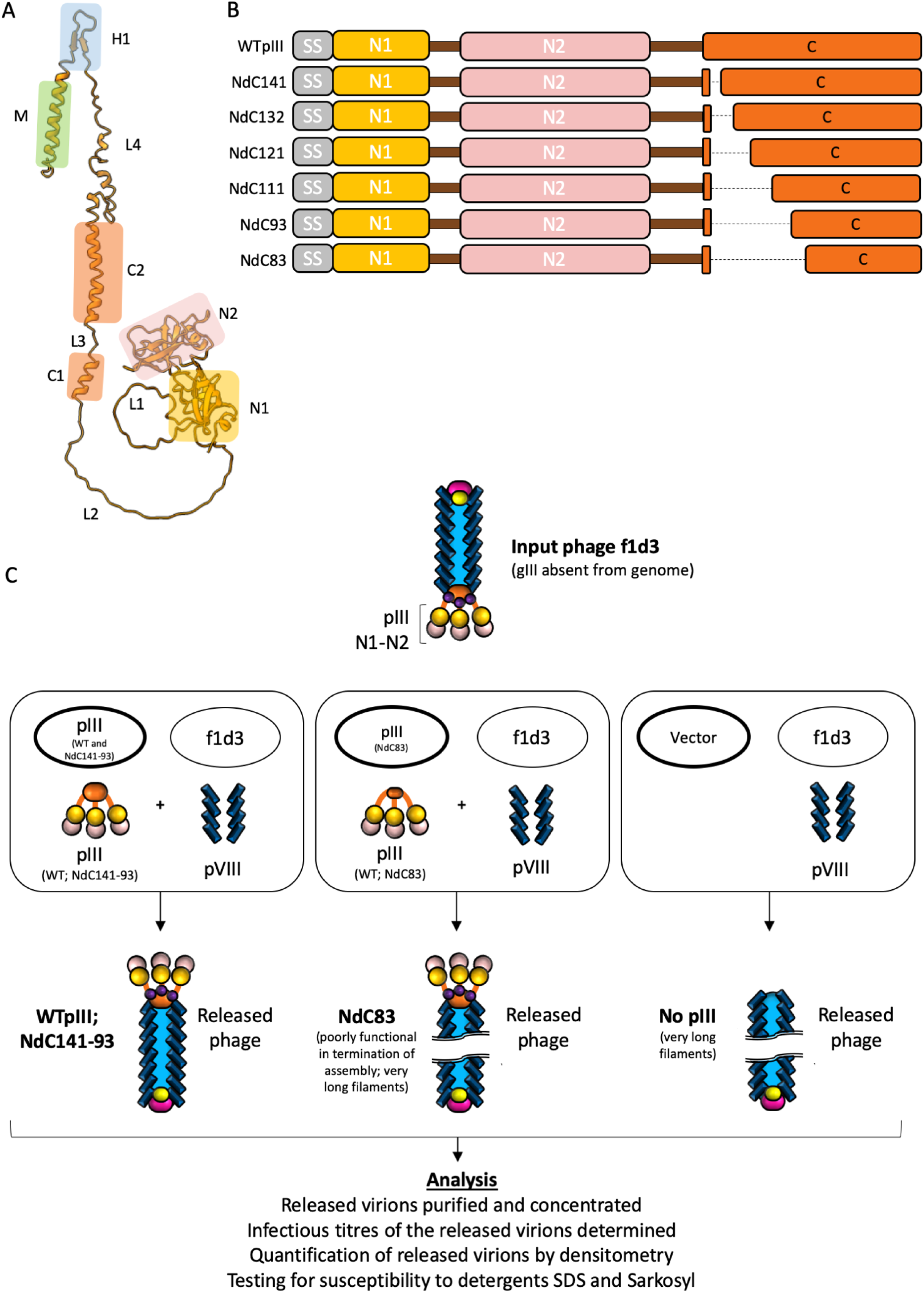
Production of f1d3 phage containing the C domain mutants. (A) Domain organisation of pIII. From the N-terminus of the protein: domain N1 (N1), glycine rich linker 1 (L1), domain N2 (N2), glycine-rich linker 2 (L2), C-terminal domain 1 (C1), linker 3 (L3), C-terminal domain 2 (C2), linker 4 (L4), β-hairpin loop (H1) and transmembrane helix (M). (B) Diagrams of wild type pIII and nested deletion (Nd) mutants (NdC141, NdC132, NdC121, NdC111, NdC93, NdC83)^40^. SS, signal sequence (grey); N1, N1 domain (light orange); N2, N2 domain (light pink); C, C-domain (dark orange). The glycine linkers are shown as dark brown boxes, and the deleted regions as black dashed lines. (C) Diagram showing experiment for production of phage containing nested deletion mutant proteins. For clarity only three pIII proteins, instead of five, are shown. Top: input f1d3 phage stock carrying a complete deletion of gIII from the genome. Full-length pIII in the phage coat is supplied *in trans* from a plasmid. This stock was produced from *E. coli* strain K1976, expressing WT pIII from plasmid pJARA112^76^. The major coat protein pVIII is shown as blue rectangles, the tip proteins pVII / pIX as pink and yellow respectively, and the tip proteins pIII / pVI as orange and purple respectively. The N1 and N2 domains of pIII are shown in light orange and light pink respectively, as per panel B. Boxed panels: host cell plasmid (bold oval) and phage combination (fine oval) used for production and the expected composite phage particle shown underneath. Left and centre box: WTpIII and NdC141-NdC83 mutants are expressed from plasmid pJARA200, under the control of a *lacUV5* promoter^40^. NdC83 is shown separately (middle box), as most of the phage particles remain attached to the host cells, with a small fraction released into the media as very long multi-length filaments, as per the negative control^17^ (right box). Right box: non-complemented f1d3 infection (using vector pBR322 that does not supply pIII).

**Supplementary Figure 16.**
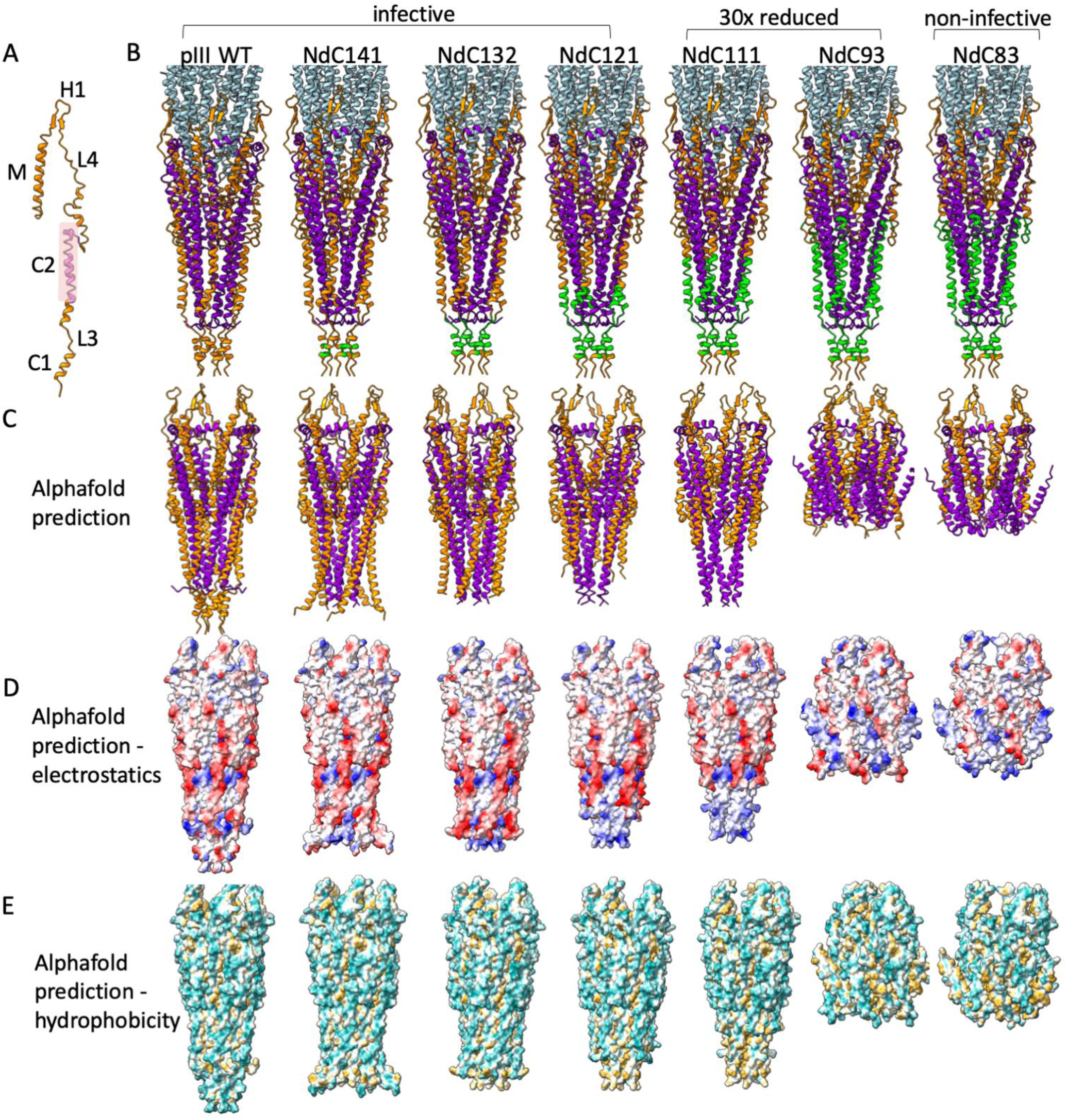
Functional analysis of the C domain in infection. (A) pIII domains, from the N-terminus of the C-domain: C-terminal domain 1 (C1), linker 3 (L3), C-terminal domain 2 (C2), linker 4 (L4), β-hairpin loop (H1) and transmembrane helix (M). The N1 and N2 domains and glycine-rich linkers are not shown. The infection-competence segment (ICS) is coloured in pink. (B) C-terminal deletion mutants are mapped onto the pointy tip structure in green - from left to right: wild type, NdC141, NdC132, NdC121, NdC111, NdC93 and NdC83. (C) Alphafold predictions of the mutation series shown as ribbons. pIII is shown in orange and pVI in purple. (D) Alphafold predictions of the mutation series shown as surface representation and coloured by electrostatic potential. Negative residues are shown in red and positive residues in blue. (E) Alphafold predictions of the mutation series shown as surface representation coloured by hydrophobicity. The most hydrophilic residues are coloured in cyan and the most hydrophobic in mustard yellow. As more pIII is removed, more of the hydrophobic helices of pVI are exposed, which begin to fold up in the mutants with the largest deletions.

**Supplementary Figure 17.**
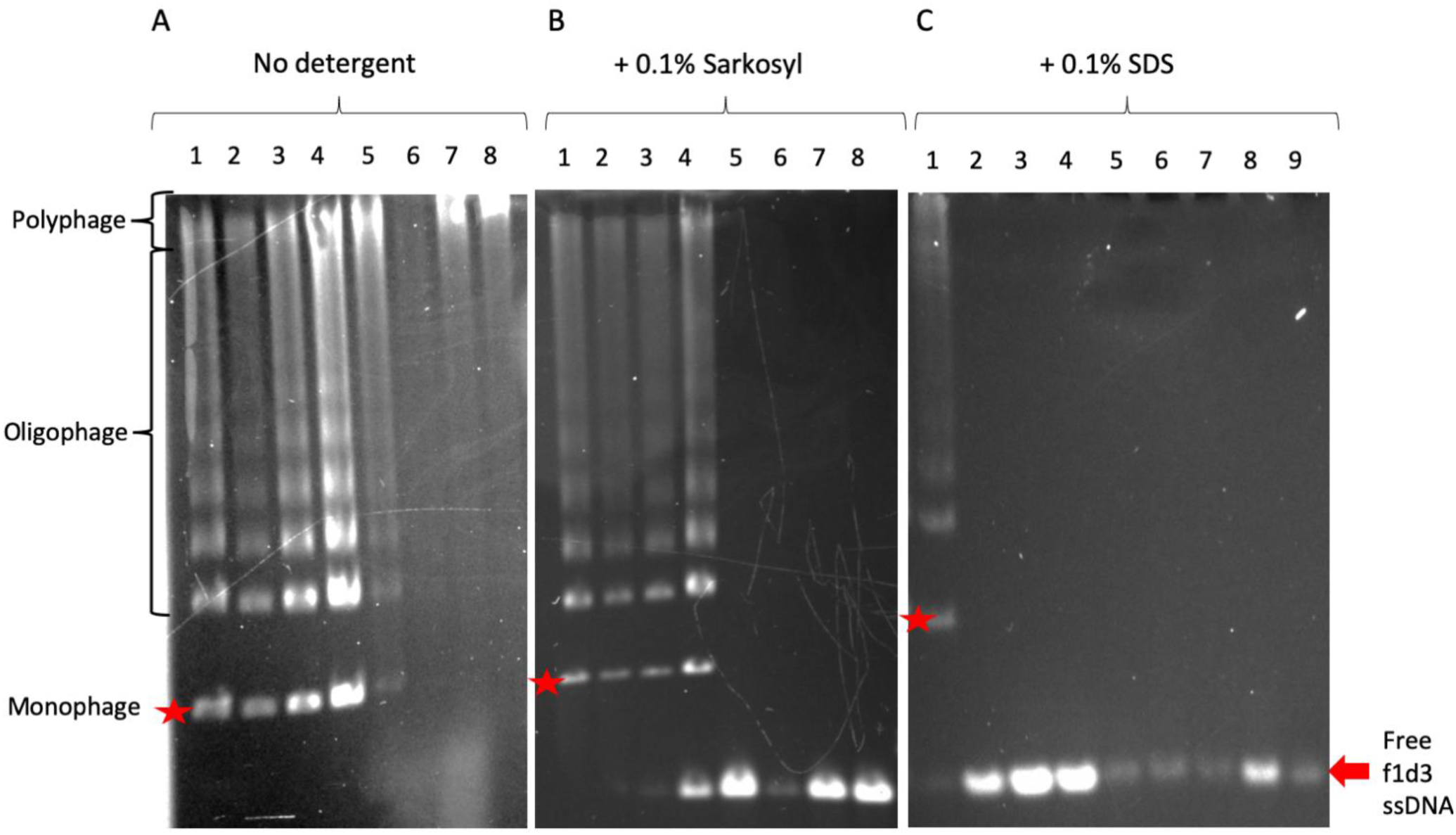
Native agarose gel electrophoresis to test the stability of phage to detergents with different properties. Panels show migration of phage and released (free) phage ssDNA in agarose gels (A) without or (B) with pre-incubation in 0.1% sarkosyl or (C) in 0.1% SDS. The position of the virions in the gel is only visible after *in situ* disassembly by soaking the gel in an alkaline solution (see Supplementary methods). Numbered lanes: Phage containing (1) WT pIII; (2) NdC141; (3) NdC132; (4) NdC121; (5) NdC111; (6) NdC93; (7) NdC83; (8) no pIII; (9) purified f1d3 ssDNA (free DNA standard). Phage containing WT pIII serves as a standard for the native virions. The f1d3 phage complemented by pIII expressed *in trans* have a wide distribution of length. Monophage (red asterisk) are phage particles containing a single f1d3 genome. Oligophage are phage particles containing two to several sequentially packaged f1d3 genomes. Polyphage are multiple-length phage particles containing sequentially packaged f1d3 genomes. Free f1d3 DNA (red block arrow) is ssDNA released from the phage during pre-incubation with detergents, or purified f1d3 ssDNA used as a standard. The analysis was performed in triplicate.

**Supplementary Figure 18.**
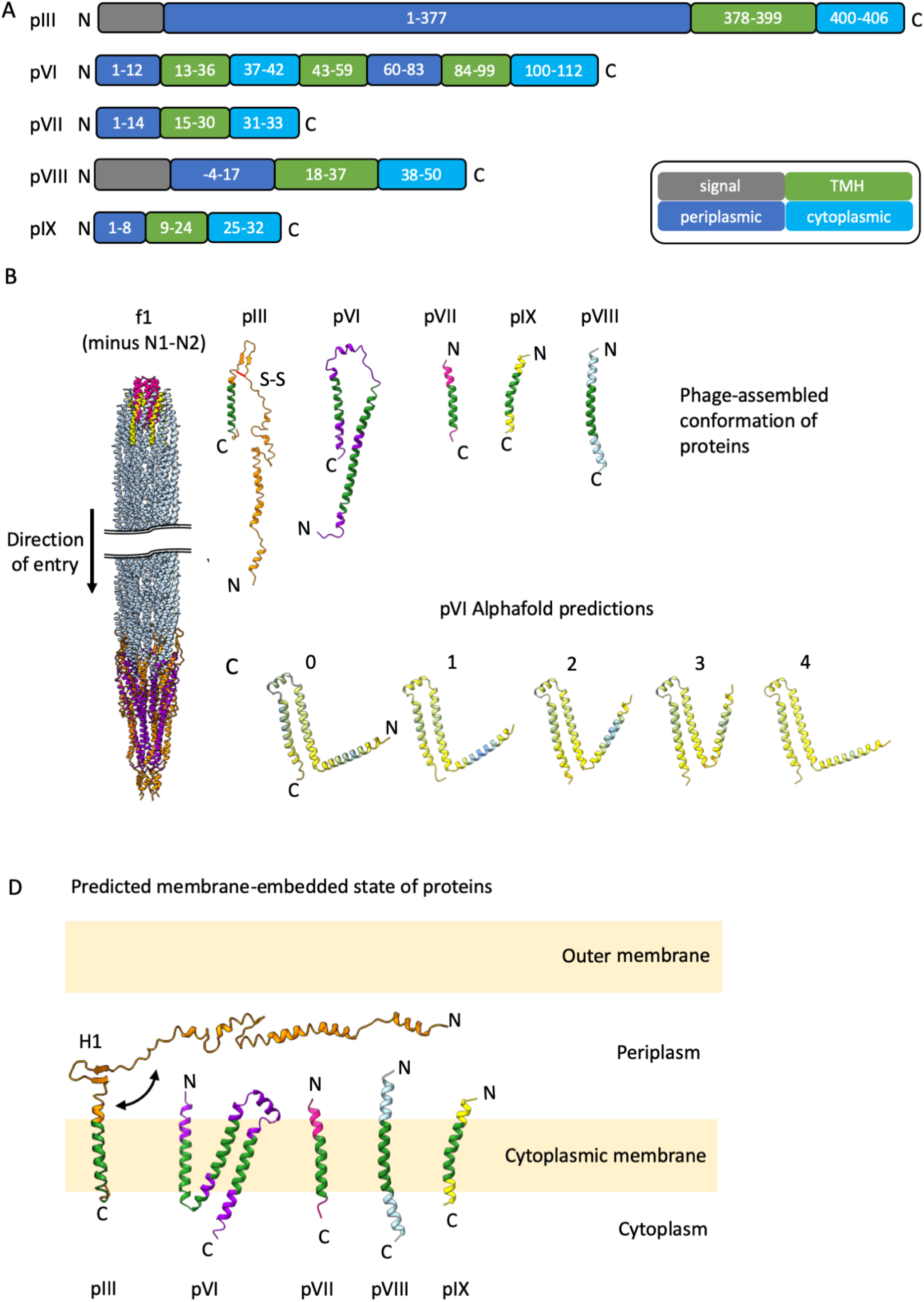
Membrane insertion of phage capsid proteins. (A) Transmembrane region predictions using MEMSAT-SVM^41^ via the PSIPRED server^77^. Signal sequences are shown in grey, transmembrane helices (TMH) in green, periplasmic regions in dark blue and cytoplasmic regions in cyan. The numbering of pIII and pVIII is for the mature proteins (excluding their signal sequences). The signal sequence predicted for pVIII varies slightly from the Uniprot entry which shows the signal sequence to consist of residues 1-23, hence the negative numbering for the periplasmic domain. (B) For clarity, a version of f1 phage without the N1-N2 domains of pIII is shown with the orientation of individual proteins in their phage-assembled conformations. The predicted transmembrane regions are coloured green. (C) Alphafold predictions of the pVI protein coloured by confidence levels with highest confidence in blue, through cornflower blue, then yellow to the lowest confidence in orange. Numbers relate to the confidence ranking from Alphafold (0 is best, 4 is worst). The N and C-termini are labelled in model 0. Interestingly, Alphafold predicts that the long N-terminal helix of pVI seen in the phage-assembled structure is folded up into two smaller helices, in agreement with the transmembrane prediction in (A). (D) Orientation of phage proteins in the cytoplasmic membrane according to MEMSAT-SVM prediction. The pIII conformation was modelled by swinging the N-terminal part of the C-domain out around the β-hairpin loop (H1). The pVI conformation is from Alphafold model 3. Proteins are coloured as above with the N and C-termini labelled.

**Supplementary Figure 19.**
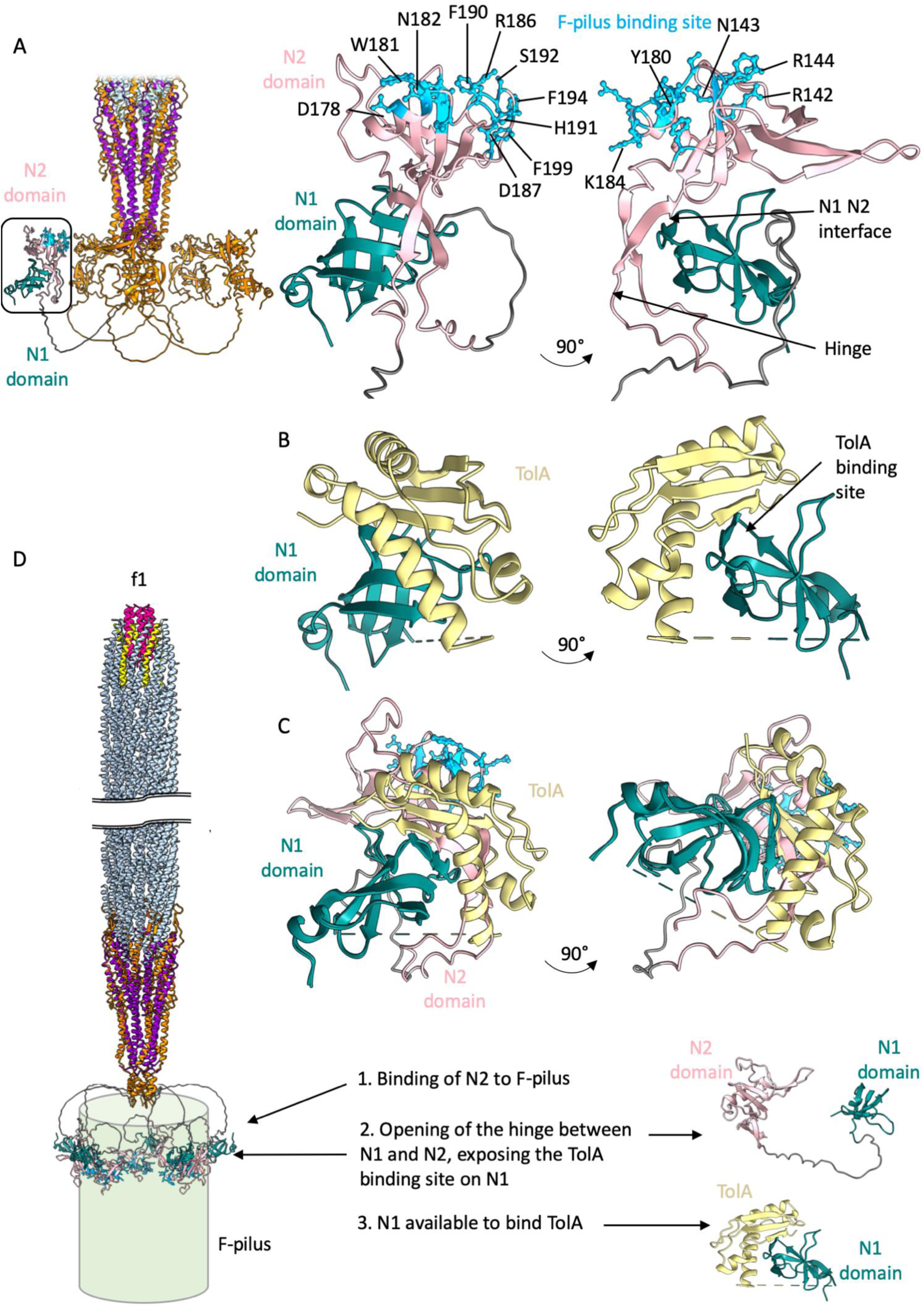
The F-pilus and TolA binding sites on pIII. (A) Overview of the pointy tip to show the N1 and N2 domains attached via flexible linkers, based on PDB 1G3P^31^. N1 is shown in teal, N2 in pink. The boxed region is shown in detail in the following panels, with the residues that make up the F-pilus binding site on N2 shown as cyan ball-and-stick (in two views 90° apart). (B) The N1 domain (teal) complexed with TolA (yellow), shown with N1 in the same orientations as in part A, based on PDB 1TOL^15^. (C) Overlay of the N1-N2 structure and N1-TolA complex in 2 views (overlaid on the N1 domains) to show that N2 and TolA share the same binding site on the N1 domain. (D) The N1-N2 domains of pIII could bind to a point on the F-pilus due to flexibility in the glycine-rich linkers between N1-N2 and the C domain of pIII. 5 copies of N2 are shown bound to the F-pilus, but it is also possible that infection occurs after a single N2 binding event. Binding of N2 to the F-pilus likely causes a conformational change in the N1-N2 domains, displacing N1 from N2 by movement of the hinge region between the two, and consequently exposing the TolA binding site on N1. The open N1-N2 structure was made by manually opening the flexible hinge region in PBD 1G3P^31^, guided by the position of N1 bound to TolA in PDB 1TOL^15^.

**Supplementary Figure 20.**
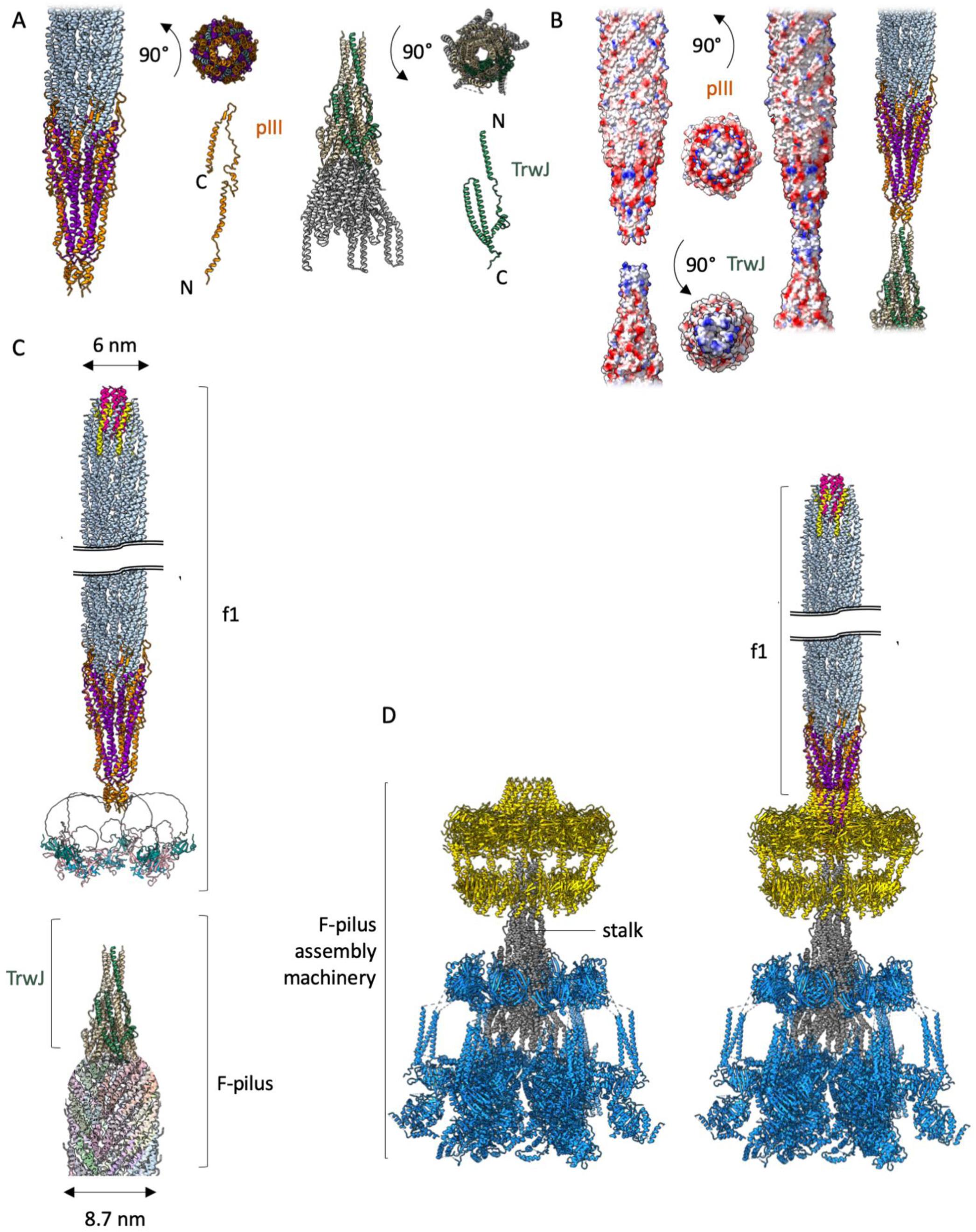
f1 binding to the F-pilus. (A) The f1-derived nanorod pointy tip and F-pilus stalk drawn to scale as cartoons, in side and top views. pIII is shown in orange and TrwJ (7O3V)^45^ in wheat, with one subunit coloured green. The remainder of the stalk is coloured in charcoal. An individual chain of pIII is shown in orange and of TrwJ in green. (B) f1 and TrwJ could interact due to opposing charge at the tip. Surface representation of both proteins, with negative residues shown in red and positive residues in blue. The proteins are shown in side and top views, and joined together as both a surface representation and ribbon diagram. pIII is shown in orange and TrwJ in wheat, with one subunit coloured green. (C) f1 phage with N1-N2 domains is drawn to scale with respect to an F-pilus (adapted from 5LER^78^, pastel colours) and TrwJ at the tip (wheat, with one subunit coloured green). (D) Left, F-pilus machinery with stalk, the tip of which is comprised of TrwJ. Outer membrane rings are shown in yellow, the stalk in charcoal, and the inner membrane complex in blue. Right, F-pilus machinery with the f1- derived nanorod.

**Supplementary Figure 21.**
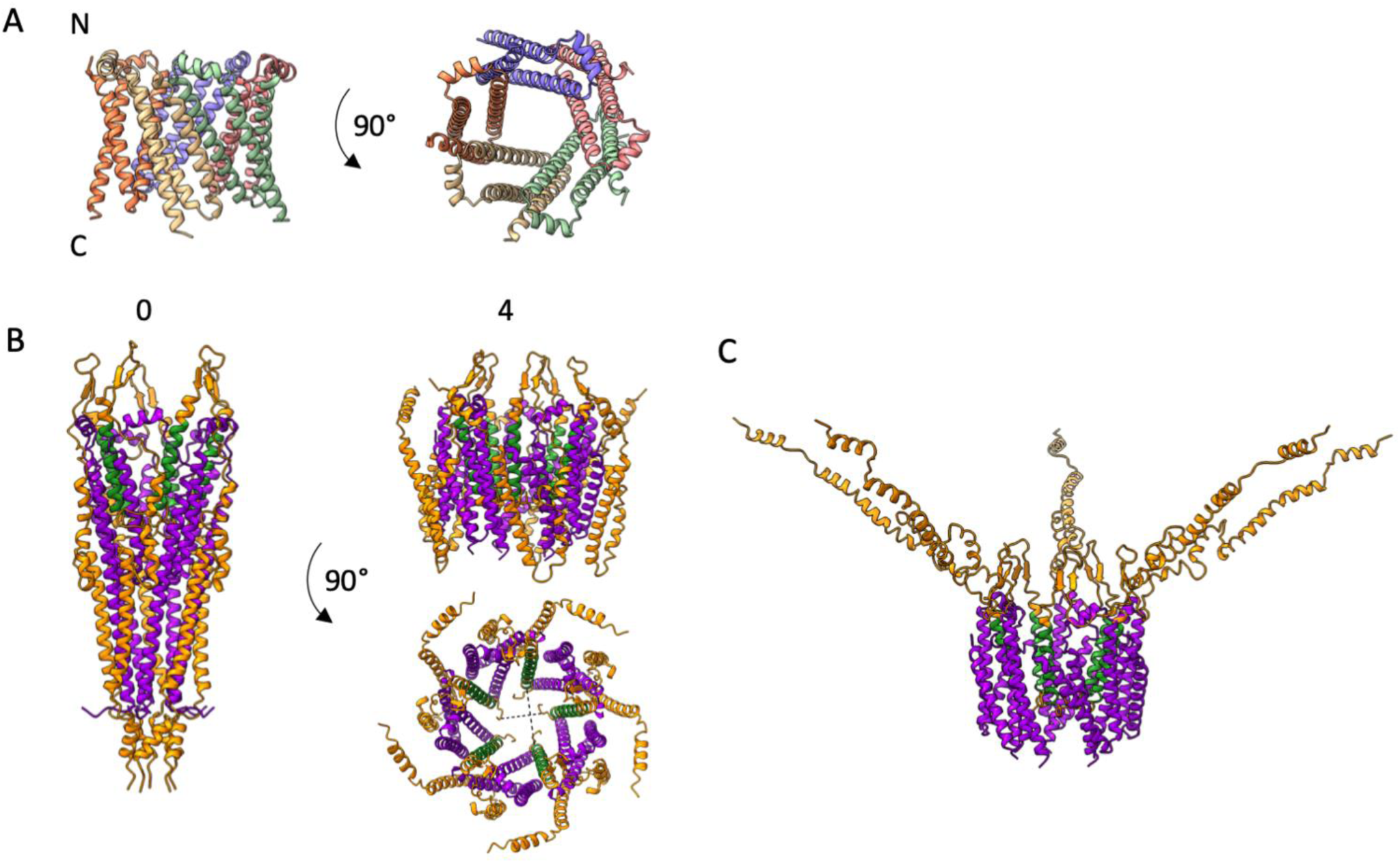
Alphafold predictions of the pVI pentamer, and pIII-pVI pentameric complex. (A) Top Alphafold prediction of the pVI pentamer coloured by chain in side and top views, depicting a pore. (B) Top Alphafold prediction (model 0) of a pentamer of the pIII C domain (orange) with a pVI pentamer (purple). This prediction is compared to the fifth Alphafold prediction (model 4), the latter being more reminiscent of how the complex might look when embedded in the cytoplasmic membrane, with the proteins in a folded up conformation. The predicted transmembrane helix of pIII is coloured green. The pointy end of the tip could peel back to allow insertion of the hydrophobic helices of pVI into the cytoplasmic membrane. A view shown looking down the channel reveals the dimensions of the pore. The diameter of the narrowest part of the open pore is between the C-terminal unstructured residues of pIII and measures 18 Å (horizontal dotted line). These unstructured regions could likely move away to allow DNA to pass through the aperture between the helices of pIII, measured at 24 Å (vertical dotted line); the diameter of a DNA double helix is 20 Å. (C) The pIII chains in the Alphafold model from part (B) were overlaid with a modelled open form of pIII, where the N-terminal part of the C-domain has been swung out around the β-hairpin loop (as described in Supplementary Fig. 18D). These helices would be free in the periplasm, in accordance with the MEMSAT-SVM predictions.

**Supplementary Figure 22.**
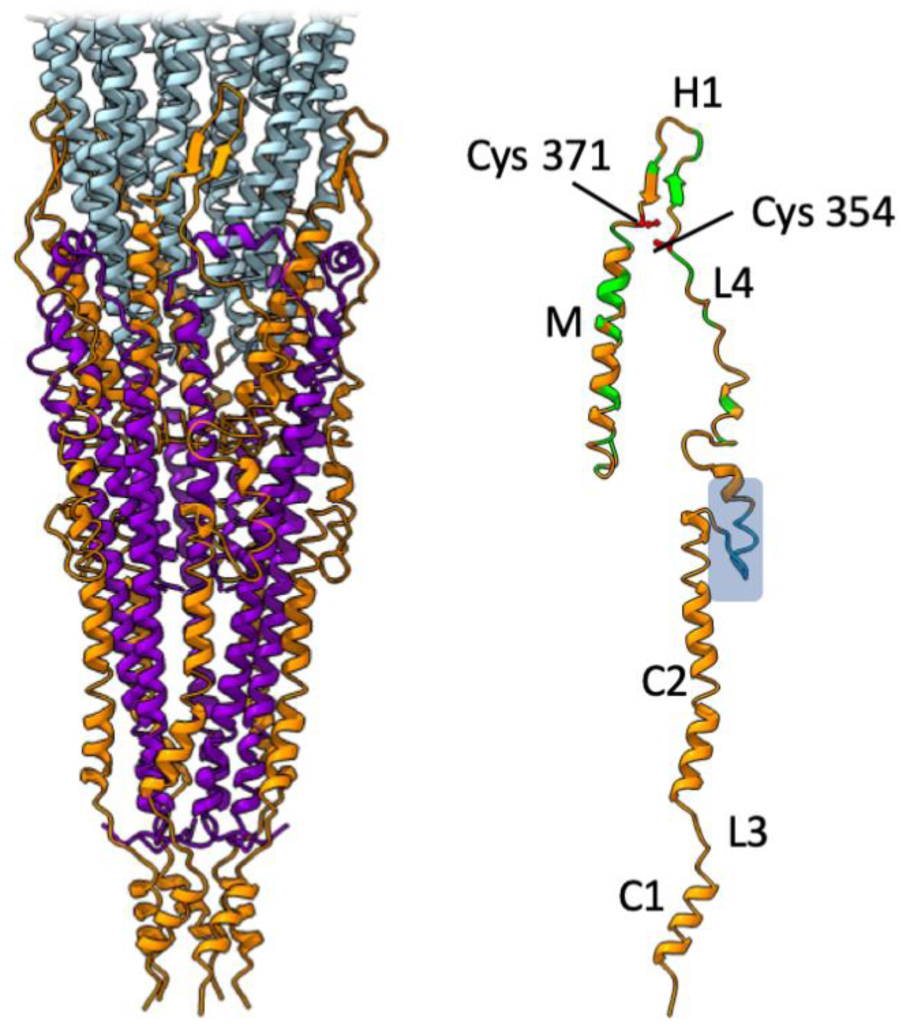
Functional analysis of the C domain in assembly and egress. pIII domains, from the N-terminus of the C-domain: C-terminal domain 1 (C1), linker 3 (L3), C-terminal domain 2 (C2), linker 4 (L4), β-hairpin loop (H1) and transmembrane helix (M). The N1 and N2 domains with glycine-rich linkers are not shown for clarity. Residues necessary for efficient incorporation of pIII into the wild-type coat^58^ have been mapped onto our structure and shown in green. Cys 354 and Cys 371 were also shown to be essential for assembly^59^ and are shown as ball-and-stick in red and labelled. The 10 amino acid long region required for termination of assembly^23^ is coloured in blue and boxed.

**Supplementary Table 1.**
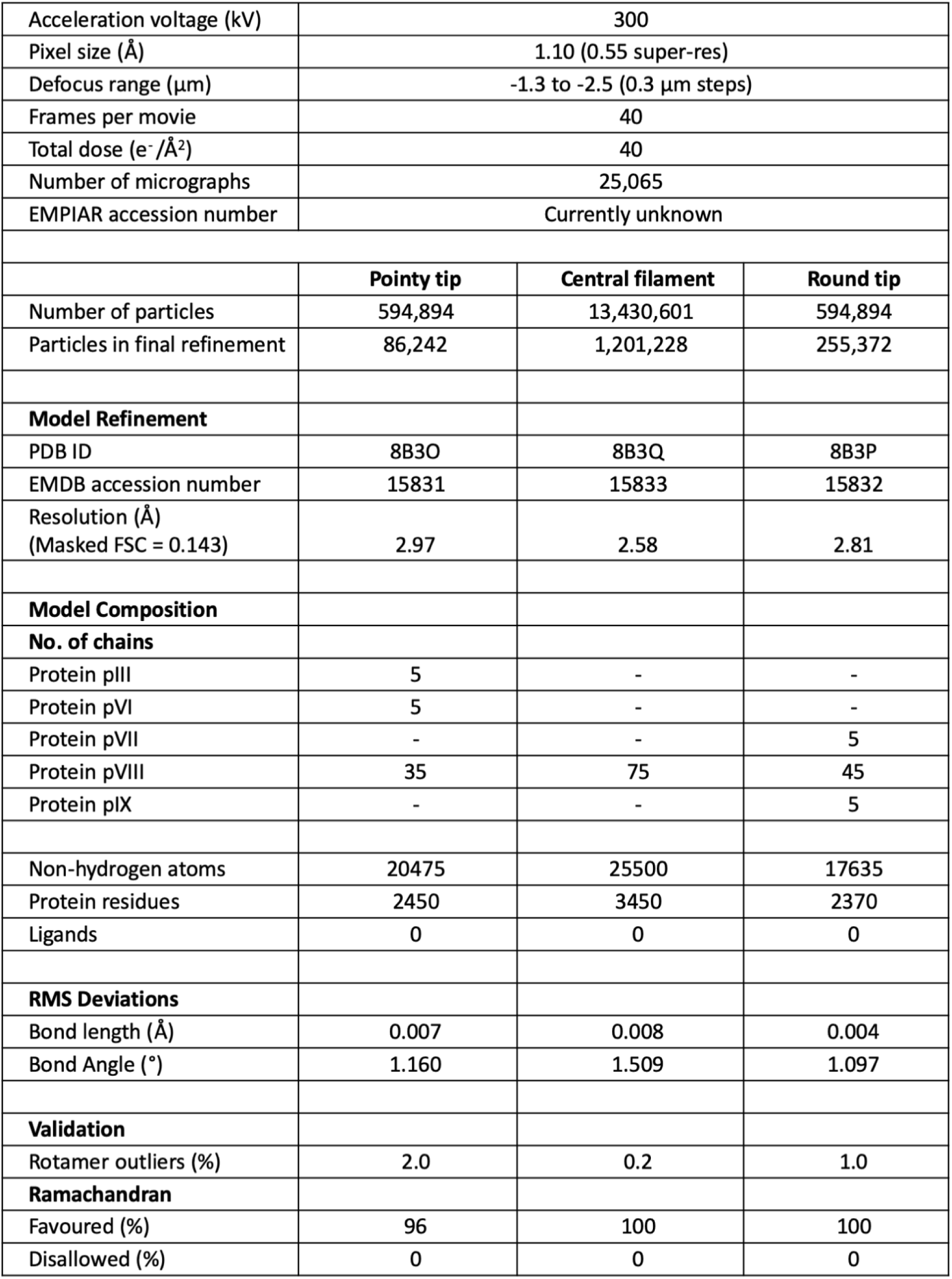
CryoEM data collection and processing details.

|  |  |  |  |
| --- | --- | --- | --- |
| Acceleration voltage (kV) | 300 |  |  |
| Pixel size (Å) | 1.10 (0.55 super-res) |  |  |
| Defocus range (µm) | -1.3 to -2.5 (0.3 µm steps) |  |  |
| Frames per movie | 40 |  |  |
| Total dose (e<sup>-</sup>/Å<sup>2</sup>) | 40 |  |  |
| Number of micrographs | 25,065 |  |  |
| EMPIAR accession number | Currently unknown |  |  |
|  | Pointy tip | Central filament | Round tip |
| Number of particles | 594,894 | 13,430,601 | 594,894 |
| Particles in final refinement | 86,242 | 1,201,228 | 255,372 |
| Model Refinement |  |  |  |
| PDB ID | 8B3O | 8B3Q | 8B3P |
| EMDB accession number | 15831 | 15833 | 15832 |
| Resolution (Å)<br>(Masked FSC = 0.143) | 2.97 | 2.58 | 2.81 |
| Model Composition |  |  |  |
| No. of chains |  |  |  |
| Protein pIII | 5 | - | - |
| Protein pVI | 5 | - | - |
| Protein pVII | - | - | 5 |
| Protein pVIII | 35 | 75 | 45 |
| Protein pIX | - | - | 5 |
| Non-hydrogen atoms | 20475 | 25500 | 17635 |
| Protein residues | 2450 | 3450 | 2370 |
| Ligands | 0 | 0 | 0 |
| RMS Deviations |  |  |  |
| Bond length (Å) | 0.007 | 0.008 | 0.004 |
| Bond Angle (°) | 1.160 | 1.509 | 1.097 |
| Validation |  |  |  |
| Rotamer outliers (%) | 2.0 | 0.2 | 1.0 |
| Ramachandran |  |  |  |
| Favoured (%) | 96 | 100 | 100 |
| Disallowed (%) | 0 | 0 | 0 |

**Supplementary Table 2.**
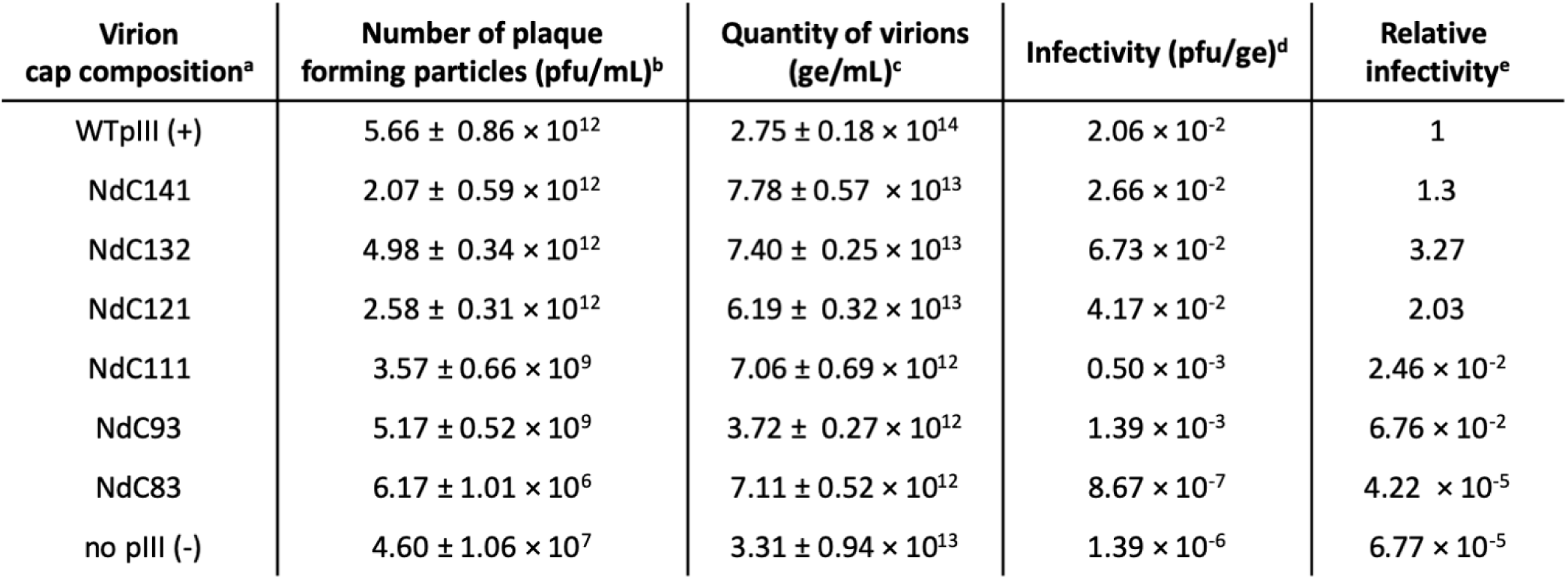
The infectivity of pIII C domain mutants. ^a^Phages were produced as described in Supplementary methods. ^b^Titre; the number of infectious phage particles per ml. ^c^Quantity of phage particles, expressed as the number of genome equivalents (ge) per ml. A genome equivalent is a measure of particle mass and is defined as a particle containing one encapsulated genome (or the portion of a multiple length particle that contains one genome). Thus, a virion particle containing ten genomes represents ten genome equivalents, as do ten phage particles containing one genome each. The number of genome equivalents was determined from agarose gel electrophoresis of phage ssDNA, released from SDS-disassembled virions^40^ as described in Supplementary methods. ^d^Infectivity of phage, expressed as a ratio of infectious titre to quantity of phage particles. ^e^Relative infectivity expressed as the ratio of infectivity of phage particles relative to those produced in the cells expressing WT pIII (positive control). Virion quantification and titration of the infectious particles was carried out in triplicate.

## Supplementary methods

### Determination of the nanorod length from micrographs of negatively stained nanorods

Purified nanorods (10 μl of 5x10^14^ particles/ml) were applied onto carbon coated 400 mesh copper grids (Electron Microscopy Sciences) for 5 min. The grid was then removed from the drop and the excess liquid was removed with filter paper before placing the grid on top of a 10 μl drop of MilliQ water for 30 sec as a washing step. Next, the grid was removed from the drop and the excess liquid was removed with filter paper. Finally, the grid was placed on top of a 10 μl drop of 2% uranyl acetate for 5 min to stain the sample, followed by the removal of the excess liquid. Images were obtained using a 120 kV Tecnai G2 Spirit BioTWIN (FEI, Czech Republic) microscope at the Manawatu Microscopy and Imaging Centre (MMIC), Palmerston North, New Zealand. Nanorod size was determined by measuring the length of 300 well-separated particles using the ImageJ^65^ measurement function.

### Production of C domain mutants

The host strains for phage production were obtained by transformation of the NdC141-NdC83 series^40^ or the WTpIII-expressing plasmids into strain TG1 (*supE Δ(hsdM-mcrB)5* (rk^-^ mk^-^ McrB^-^) *thi Δ(lac-proAB*) F’ [*traD36 lacI^q^Δ(lacZ)M15 proA^+^B^+^*]). Transformants were selected for on 2xYT plates containing 100 µg/ml Amp and were colony-purified. Single colonies were used to inoculate overnight cultures for phage production. To produce phages containing WT pIII or mutants with internal C domain deletions, exponentially growing cultures were infected with phage f1d3 at a multiplicity of infection of 100 phage per cell, for 1 hour at 37 °C. Infected cells were then separated from unabsorbed phage by centrifugation (7000 x *g* for 10 minutes at 37 °C), resuspended in fresh medium containing 0.1 mM IPTG and incubated for a further period of 4 h to allow phage or phagemid particle production. Following incubation, the cells were removed by centrifugation (7000 x *g* for 20 min at 30 °C) and the supernatants containing the released phage were collected for further analyses. Phage in the supernatants was concentrated by two rounds of standard PEG precipitation [PEG8000 (5 % w/v) and NaCl (0.5 M)].

### Infectivity of C domain mutants

Briefly, virions were disassembled by incubation in SDS-containing buffer (1% SDS, 1× TAE Buffer 5% Glycerol, 0.25% BPB) at 70°C for 20 min to release the phage ssDNA and the samples were subjected to agarose gel electrophoresis. On completion of electrophoresis, phage ssDNA was stained with ethidium bromide and quantified by densitometry. Since the amount of ssDNA in a band is not linearly proportional to the intensity of the fluorescence, every gel contained a set of twofold serial dilutions of f1d3 ssDNA standard of known concentration, used for calibration. Quantity of phage particles was expressed as the number of genome equivalents (ge) per ml. A genome equivalent is a measure of particle mass and is defined as a particle containing one encapsulated genome (or the portion of a multiple length particle that contains one genome). Thus, a virion particle containing ten genomes represents ten genome equivalents, as do ten phage particles containing one genome each.

Infectivity of phage was expressed as a ratio of infectious titre to quantity of phage particles. Infectious titre was determined by titration of samples on the strain TG1 as a host. For each sample the titre was determined from counts on three plates, each containing between 100 and 300 plaques. Relative infectivity was expressed as the ratio of infectivity of phage particles relative to those produced in the cells expressing WT pIII (positive control).

### Testing detergent sensitivity of phage

Samples were mixed with DNA loading buffer (1× TAE, 5% Glycerol, 0.25% BPB) and loaded onto 0.5% agarose gels. Different detergents were added to the loading buffer and the sample incubated at room temperature for ten minutes prior to loading. The phage particles were separated by electrophoresis at 1.3 V/cm overnight. After electrophoresis, free DNA released from phage by detergent pre-treatment was detected by staining the gel in ethidium bromide. To detect the phage, the gel was soaked in 0.2 M NaOH for 1 hour to disassemble the virions, followed by neutralisation in 0.45 M Tris pH 7.1. The exposed phage ssDNA was then visualised by staining the gel again with ethidium bromide^40^.

